# Scalable Multi-omics Diagonal Integration from Self-Mapping

**DOI:** 10.64898/2026.11.10.765642

**Authors:** Chuan He, Leying Guan

## Abstract

Diagonal integration seeks to align single-cell multi-omics datasets in which distinct modalities, such as RNA and protein, are measured in different cells. Unlike horizontal integration, these datasets lack direct cell–cell correspondences and complete feature mappings across modalities. Leading methods for weak-linkage integration often infer anchor cells between modalities by searching for cross-modal similarities through predefined feature links, then use these anchors to guide alignment. Their required matching or anchor refinement can become computationally burdensome at atlas scale. Here, we introduce a novel self-mapping technique, a mechanism that aligns modalities by mapping each computationally decoded cross-modality omics profile prediction to its own observed inputs. We implement this strategy in SCALEMAP, a scalable, modality-agnostic method for diagonal integration. Across transcriptomic, epigenomic and proteomic benchmarks, SCALEMAP matches or outperforms leading alternatives while requiring substantially less runtime and memory. In large-scale experiments, SCALEMAP integrates more than 1.5 million multi-omic profiles in less than 5 minutes with under 20 GB peak memory, which enables rapid synthesis of massive single-cell multi-omics resources into unified biological representations.

## 1 Introduction

Single-cell technologies now measure cellular states through transcriptomic, epigenomic and proteomic readouts, providing complementary views of cell identity and regulation [1–6]. Assays such as CITE-seq and TEA-seq can profile multiple modalities in the same cell and therefore provide direct cross-modality correspondence [5, 7]. However, many large and biologically valuable resources remain single-modality, spatial, targeted, or otherwise generated from distinct cell populations [6, 8–11]. Integrating these resources requires methods that can align cells across different feature spaces without assuming that the same cell has been measured in each modality.

This setting is known as diagonal integration [12]. Unlike horizontal integration, where datasets share a measured feature space, diagonal integration lacks both direct cell–cell correspondence and complete feature correspondence across modalities. A simple strategy is to construct a surrogate shared feature space from known biological links, such as gene expression and gene-activity scores for RNA–ATAC integration, or protein abundance and expression of the corresponding coding genes for RNA–protein integration. Once both modalities are represented by these linked features, conventional shared-feature integration methods, including anchor-based matching, or latent-space batch correction, can be applied by treating modality as a dataset or batch label [13–16]. However, these methods perform the initial alignment using only the surrogate shared features and therefore depend on those features providing a sufficiently informative and comparable representation of cellular state across modalities. In weak-linkage settings, the available links are often sparse, noisy or biologically indirect, so cross-modality cell similarity computed from these features may provide an unstable alignment signal. RNA–protein integration is a representative example: transcript and protein abundance can differ because of post-transcriptional regulation and related processes [17–20], and targeted protein panels often contain far fewer features than transcriptomic profiles.

Several tailored approaches have been developed for diagonal integration under limited feature correspondence. Geometry-based methods, including Gromov–Wasserstein optimal transport approaches such as SCOT and Pamona [21, 22], avoid direct cross-modality feature comparison by matching within-domain cell–cell geometry across modalities. These methods do not require predefined feature correspondences, but they are computationally demanding and intrinsically underdetermined: similar within-domain geometry does not uniquely specify which cells should correspond across modalities. Recent state-of-the-art weak-linkage methods therefore focus on learning or refining cross-modality cell relationships from limited feature correspondence. MaxFuse uses fuzzy smoothing and iterative coembedding to refine cross-modality cell matching [23]. bindSC uses CCA-based iterative bi-order integration to jointly learn cross-modal cell correspondences and cross-modal feature correspondences [24]. scMODAL uses linked features to construct minibatch mutual-nearest-neighbor anchors that guide deep latent-space alignment [25]. These methods improve alignment under weak linkage, but they still require explicit inference of relationships between cells from different modalities. This step can be computationally costly as the number of cells increases. Minibatch anchor construction reduces the cost of full-dataset anchor search, but reliable anchor construction still requires a sufficiently large cross-modality candidate set, which limits how far the minibatch size can be reduced without compromising the alignment signal.

Here we introduce SCALEMAP, a deep diagonal integration method that uses feature correspondence through a novel cell-specific self-mapping mechanism. Rather than using feature links to search for anchor cells, SCALEMAP uses them to map its own cross-modal translated profile to the observed inputs. This self-mapping objective gives every cell its own positive alignment target, which avoids cross-modal cross-modal cell-cell anchor search while retaining biologically grounded feature correspondence. We evaluate SCALEMAP in RNA–protein diagonal integration tasks where shared feature information is limited. Across these benchmarks, SCALEMAP provides competitive or improved integration quality across biological conservation, batch mixing and modality alignment metrics, and cross-modality protein prediction while requiring substantially less runtime and memory than leading alternatives. In large-scale experiments, SCALEMAP integrates more than 1.5 million single-cell mutl-omics profiles in less than 5 minutes with under 20 GB peak memory. We further evaluate SCALEMAP in RNA–ATAC integration, where more cross-modality feature links can be constructed but the correspondence remains imperfect, and find that SCALEMAP remains competitive or top-performing across RNA–ATAC benchmarks while retaining low runtime and memory use.

## 2 Results

### 2.1 Method Overview

Diagonal integration requires an alignment signal between modalities observed in different cells. SCALEMAP defines this signal through self-mapping. For each input cell, the model translates the observed profile into the other modality and compares the translated linked features with a linked-feature representation derived from the same input cell through predefined feature correspondence. Anchor-based workflows use feature correspondence to search for positive cross-modality cell relationships before alignment; SCALEMAP uses the feature-correspondence table after cross-modal decoding to define a cell-specific translation target.

SCALEMAP implements self-mapping within a well-established multi-input, multi-output autoen-coder (Fig. 1a). For a cell profile 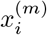 from modality *m* ∈ {1, 2}, the encoder path *E*_*m*_ maps the input into a shared latent code 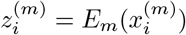. Each modality has its own decoder *G*_*m*_. From 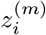, decoder *Gm* reconstructs the input modality, and decoder *G*_*¬m*_ generates a translated profile in the other modality, 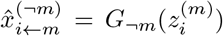, which can be interpreted as the predicted “counter-factual” for the unseen modal for cell *i*.

**Fig. 1.**
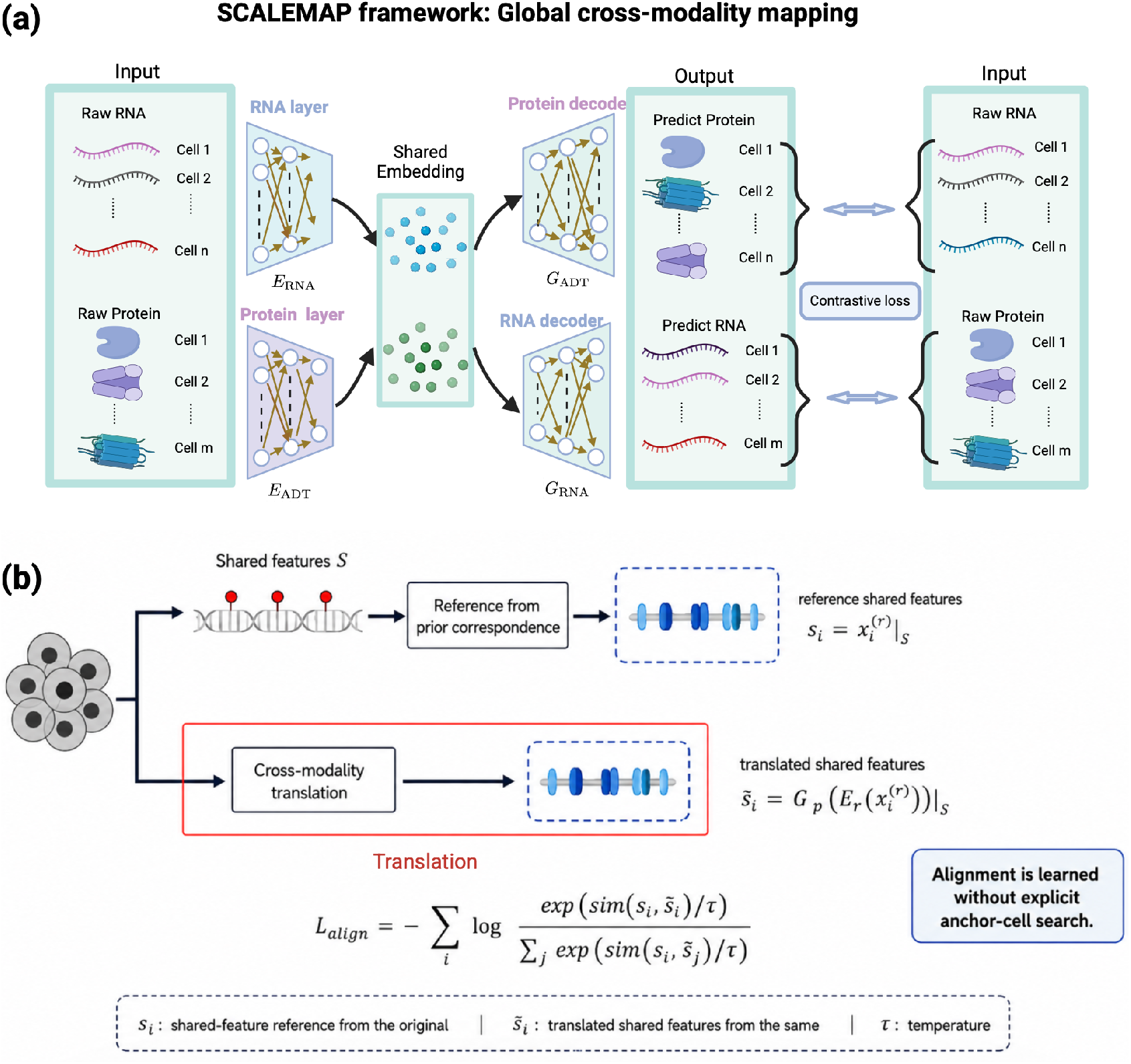
Overview of the SCALEMAP framework and comparison with anchor-based integration strategies. (a) SCALEMAP projects RNA and protein data into a shared latent space using modality-specific encoders. The latent representation supports bidirectional cross-modality translation through modality-specific decoders, enabling RNA-to-protein and protein-to-RNA prediction. (b) SCALEMAP uses shared biological feature correspondences to define reference shared features and trains cross-modality translation to match the corresponding translated shared features through a mini-batch contrastive loss.

The self-mapping target is defined in an ordered linked-feature comparison space (Fig. 1b). Let 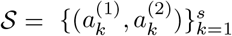 denote an ordered set of biologically linked feature pairs between the two modalities, and let *P*_*m*_ extract the modality-*m* features corresponding to these links. For a cell observed in modality *m*, SCALEMAP extracts the linked-feature representation 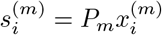 from the observed input. In RNA– protein integration, the linked pairs can be measured proteins and the genes encoding those proteins.

In RNA–ATAC integration, they can be gene expression and gene-activity values defined through gene-level correspondence. SCALEMAP then applies self-mapping to the translated profile from the same input cell. It decodes 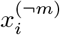, the observation from the other modality, into modality *m* and extracts the corresponding linked features in the same ordered comparison space, 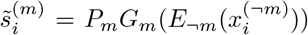. Both 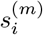 and 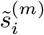 are represented in the same ordered linked-feature space after modality-specific preprocessing, so the contrastive objective compares corresponding biological links rather than treating raw measurements from different assays as identical quantities. The self-mapping objective uses 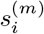 and 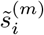 as the positive pair for cell *i*. In the minibatch contrastive loss, this same-input-cell pair is encouraged to score higher than pairs formed with translated linked-feature vectors from other cells. This contrastive objective supplies diagonal supervision without identifying matched cells from different modalities.

Self-mapping is optimized jointly with the objectives that learn the shared representation and translation functions (see Methods). These terms keep latent codes informative for the observed assays and reduce modality-driven separation in the shared space. After optimization, the shared latent codes 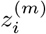 are used as the integrated cell embedding and model-specific translations.

### 2.2 SCALEMAP enables efficient diagonal integration of atlas-scale data

We assessed the scalability of SCALEMAP against four state-of-the-art multi-omics integration methods, including scMODAL, GLUE, BindSC and MaxFuse, using the recently published COMBAT atlas-scale CITE-seq dataset, which contains paired RNA and antibody-derived tag (ADT) measurements from more than 830,000 cells [26] (Supplementary Section 1). We generated six datasets comprising 2%, 5%, 10%, 20%, 50% and 100% of the full dataset. All experiments were performed on a 32-core Intel CPU, with limits of 12h for runtime and 100 GiB for memory. Across increasing dataset sizes, SCALEMAP exhibited an approximately constant runtime and near-linear memory scaling (Fig. 2a,b). SCALEMAP had the shortest runtime at every scale and integrated the full dataset in 3.3 min. By contrast, competing methods incurred substantially greater computational costs or failed to scale to the larger datasets. BindSC exceeded the memory limit at moderate dataset sizes, whereas the runtimes of MaxFuse and GLUE increased rapidly and ultimately exceeded the 12-h limit. Although scMODAL completed the analysis of the full dataset, it required 87.5 min under its default settings. These results demonstrate the computational efficiency of SCALEMAP for atlas-scale data integration.

**Fig. 2.**
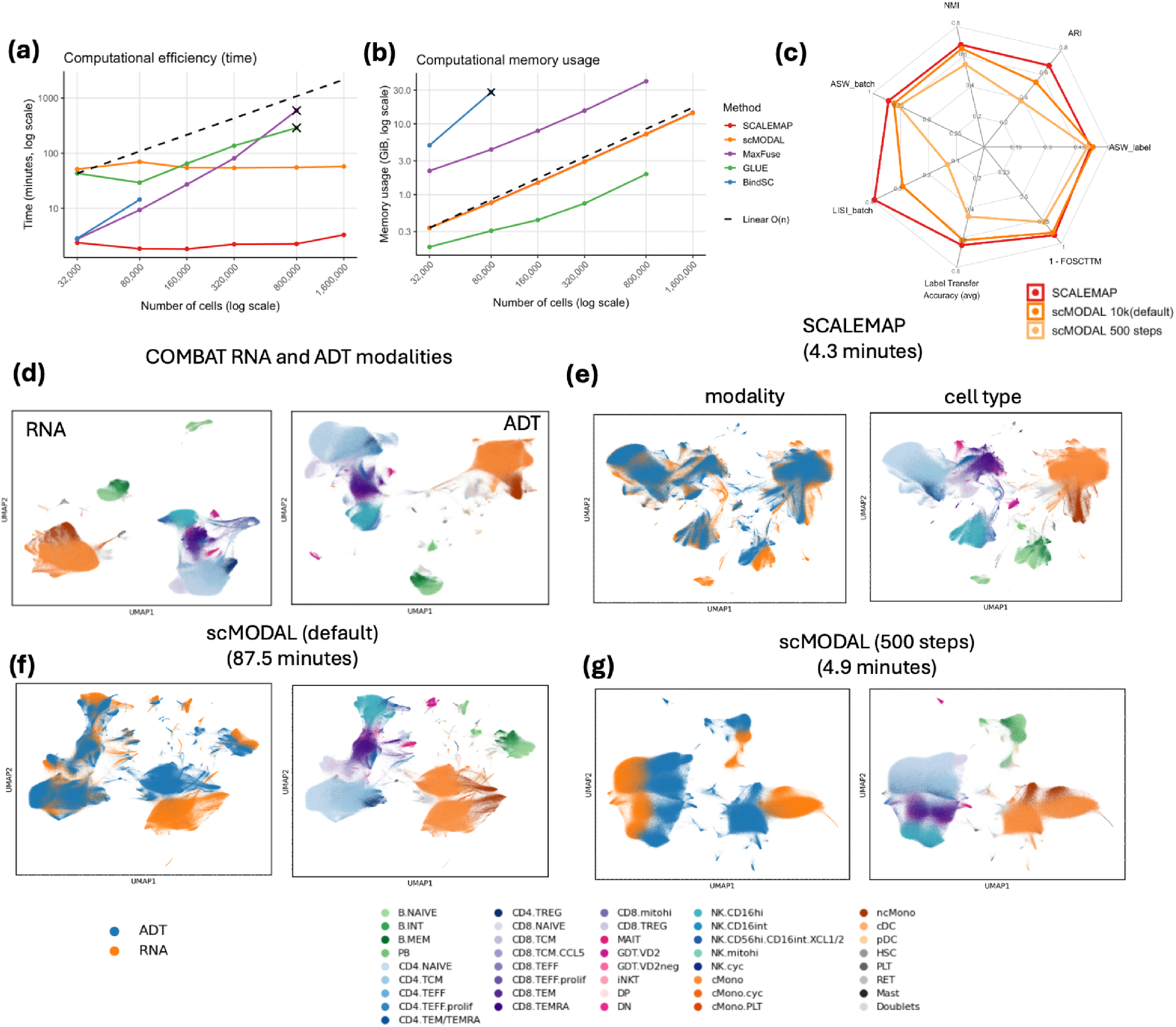
Scalability and integration performance on the large-scale COMBAT dataset. (a-b) Runtime and memory comparison across increasing dataset sizes. Several methods fail to complete at larger scales due to time or memory limits (indicated by X). (c) Radar plot of integration metrics on the full COMBAT dataset. (d-g) UMAP visualization of raw RNA and ADT modalities, SCALEMAP, scMODAL full, and scMODAL 500 epochs integrated embedding colored by batch and cell type.

SCALEMAP also outperformed scMODAL in diagonal integration across three complementary aspects of performance (Fig. 2c–f): biological conservation, quantified using label-based average silhouette width (ASW), adjusted Rand index (ARI) and normalized mutual information (NMI); batch mixing, quantified using batch ASW and batch local inverse Simpson’s index (LISI); and cross-modality matching, quantified using label-transfer accuracy and one minus the fraction of samples closer than the true match (1-FOSCTTM) (Supplementary Section 3). Relative to scMODAL, SCALEMAP achieved a 25.5% higher ARI, a 34.6% higher batch LISI score and a 23.1% higher 1-FOSCTTM score. Thus, SCALEMAP combined improved computational scalability with strong integration accuracy.

To compare the methods at similar runtimes, we reduced scMODAL training from the default 10,000 steps to 500 steps. Under this setting, scMODAL and SCALEMAP required 4.9 min and 4.3 min, respectively. However, reducing the number of scMODAL training steps substantially impaired integration performance (Fig. 2c,g). Compared with reduced-step scMODAL, SCALEMAP achieved a 76.4% higher ARI, a 199.6% higher batch LISI score and a 60.9% higher 1-FOSCTTM score. Consistent with these quantitative results, UMAP visualizations of the integrated embeddings showed clearer separation of B-cell and monocyte subtypes with SCALEMAP, together with greater co-localization of the RNA and ADT modalities (Fig. 2e,g).

### 2.3 SCALEMAP is computationally efficient while maintaining leading-tier RNA–protein integration performance

We benchmarked SCALEMAP on six RNA–protein diagonal integration tasks spanning CITE-seq, TEA-seq and ASAP-seq datasets, including Bone Marrow CITE-seq [13], PBMC TEA-seq [7], a COMBAT CITE-seq subset [26], PBMC ASAPseqCtrl [27], PBMC ASAPseqStim [27] and PBMC CITE-seq [28] (see Supplementary Section 1). These datasets differ in tissue, cell number, technology, feature dimension and the number of linked RNA–protein features, ranging from limited-linkage settings such as Bone Marrow CITE-seq to datasets with larger protein panels (Fig. 3a). For each dataset, paired RNA and protein measurements were split into modality-specific inputs to create an unpaired diagonal integration task while retaining the matched measurements for evaluation.

**Fig. 3.**
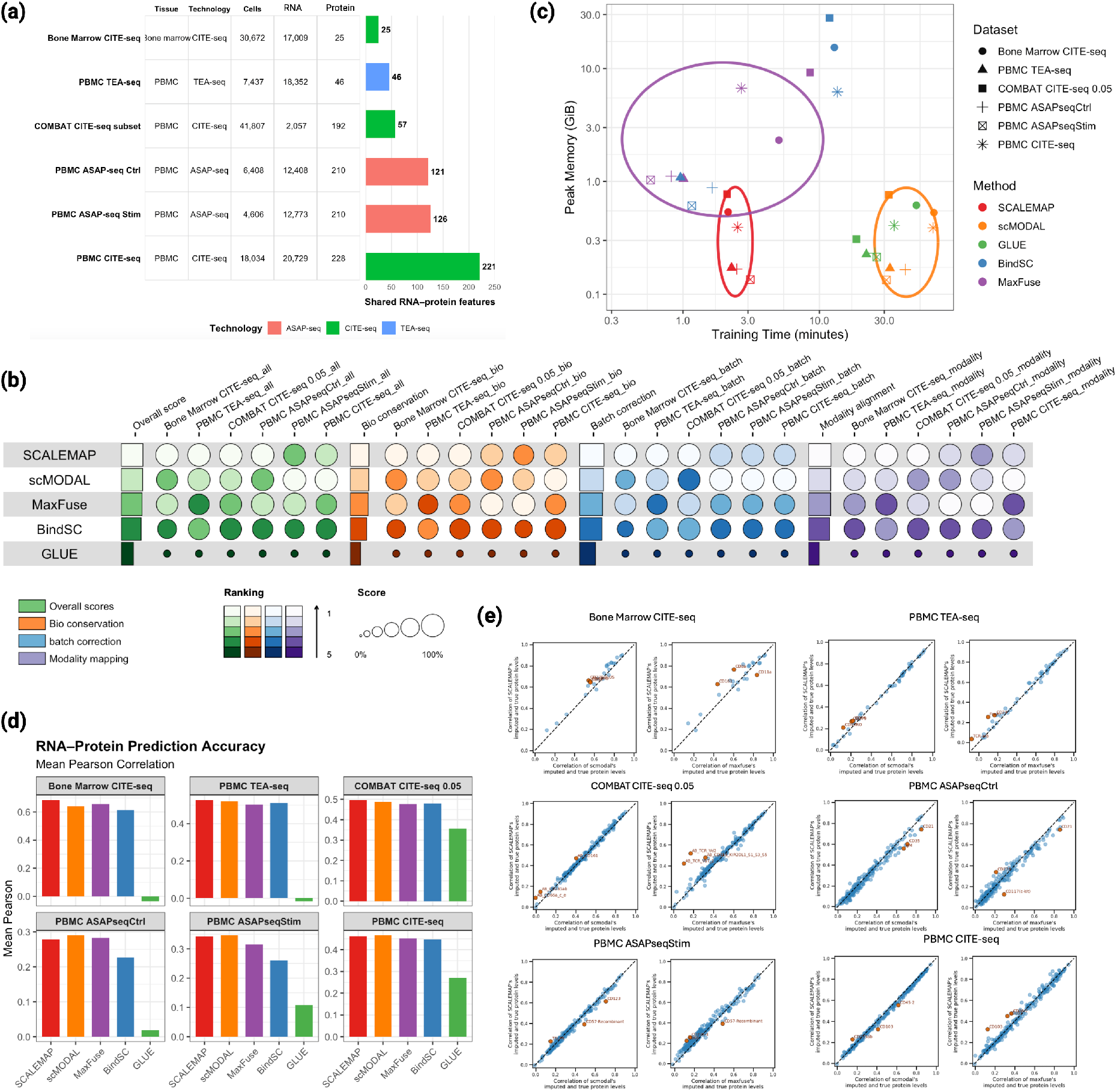
Comprehensive evaluation of integration performance, efficiency, and cross-modality prediction. (a) Overview of the six RNA–protein benchmark datasets, showing tissue source, assay technology, number of cells, RNA and ADT feature dimensions, and the number of shared RNA–protein correspondence features. Datasets are ordered by the increasing number of shared features. (b) Bubble plot summarizing integration performance across six datasets. Circle size represents the magnitude of the score (linearly scaled), and color indicates metric category: green for overall score, orange for biological conservation, blue for batch correction, and purple for modality alignment. Methods are ranked based on their overall score. (c) Runtime versus peak memory usage across datasets. Each point corresponds to a method–dataset pair, where color denotes the integration method and marker shape represents the dataset type. (d) RNA-to-protein prediction accuracy measured by mean Pearson correlation across proteins. Colors correspond to different methods with the same color code as (b). (e) Pairwise comparison of protein prediction performance between SCALEMAP and the best competing method. Each point represents a protein, comparing correlation coefficients from SCALEMAP (y-axis) against the top-performing method (x-axis). The dashed line indicates equal performance. The top three proteins with the largest differences are highlighted.

We first compared integration quality using biological conservation, batch correction and modality mapping, with these category scores summarized by an overall score. Across the six datasets, SCALEMAP achieved the highest overall performance and was competitive across all three metric categories (Fig. 3b), followed by scMODAL, MaxFuse and BindSC. The average metric differences across six datasets between the top four methods were generally modest. The greatest performance decline relative to SCALEMAP was observed for BindSC, whose biological conservation and batch correction scores were, on average, 11.3% and 8.0% lower, respectively. Across individual datasets, the reductions comparing BindSC and SCALEMAP ranged from 7.5% to 14.3% for biological conservation and from 1.8% to 16.0% for batch correction. GLUE showed substantially lower scores across datasets and metric categories than the other four methods, with overall scores approximately 40–45% lower than those of SCALEMAP across datasets. The clearest separation among top-tier methods was computational efficiency. Across datasets, SCALEMAP occupied the low-runtime and low-memory region of the time–memory plot (Fig. 3c); in comparison, scMODAL required substantially longer runtime, while MaxFuse and BindSC used much more memory. Thus, SCALEMAP achieves leading-tier integration quality while requiring substantially lower computational cost.

We next evaluated whether these quantitative differences in integration quality translated into differences in RNA-to-protein prediction accuracy using 10 nearest neighbor predictors (see Methods). The predicted protein values were compared with the ground-truth protein measurements, and performance was summarized as the mean Pearson correlation across proteins within each dataset (Fig. 3d). The prediction results were consistent with the overall integration benchmark, with SCALEMAP, scMODAL and MaxFuse forming the leading group across datasets, followed by BindSC, whereas GLUE performed substantially worse than the other methods. Overall, SCALEMAP and scMODAL ranked first and second, respectively, while MaxFuse remained competitive on several datasets. Protein-level pairwise comparisons further showed that SCALEMAP and scMODAL produced more similar prediction qualities across proteins (Fig. 3e and Fig. <u>S1</u>), including on datasets for which their aggregate performance differed only modestly. In Bone Marrow CITE-seq, where linked RNA–protein information was most limited, SCALEMAP achieved comparable or slightly better prediction accuracy than scMODAL for most proteins, with larger differences observed for only a small subset of proteins. Comparisons with MaxFuse showed greater protein- and dataset-specific divergence, indicating that MaxFuse produced a more distinct embedding-based prediction structure. The results remain similar when replacing nearest-neighbor predictors with random forest predictors for different methods (Fig. S2).

Together, these results show that SCALEMAP achieves prediction performance comparable to that of the strongest alternative methods while retaining a substantially lower runtime and memory profile.

### 2.4 SCALEMAP balances modality co-localization and sub-population resolution in limited-linkage RNA–protein datasets

We examined Bone Marrow CITE-seq and the COMBAT CITE-seq subset, the two RNA–protein benchmarks with the most limited feature linkage, to assess how the aggregate scores translate into embedding structure and marker-level prediction in detail.

On bone marrow CITE-seq, SCALEMAP achieved the strongest modality mixing among the top three leading methods, improving iLISI (batch) by 56.0% over MaxFuse and 150.3% over scMODAL (Fig. 4a,b). The UMAP is consistent with this: RNA and ADT cells co-localize broadly within annotated cell-state regions under SCALEMAP, whereas MaxFuse and scMODAL retain modality-stratified neighborhoods for several populations. This did not lead to a sacrifice in other aspects, with SCALEMAP scored 5.1% and 6.2% higher on biological conservation, and 4.5% and 4.4% higher on modality alignment, than MaxFuse and scMODAL respectively. Sub-populations within both the lymphoid and progenitor/dendritic cell compartments also showed better cross-modality label correspondence using SCALEMAP, as supported by local cross-modality kNN analyses (see Methods): SCALEMAP achieved the highest macro opposite-modality label agreement in both small lymphoid and DC/progenitor compartments; it also achieved the highest hit@k in small lymphoid populations, although MaxFuse scored slightly higher in DC/progenitors (Fig. S3a-c).

**Fig. 4.**
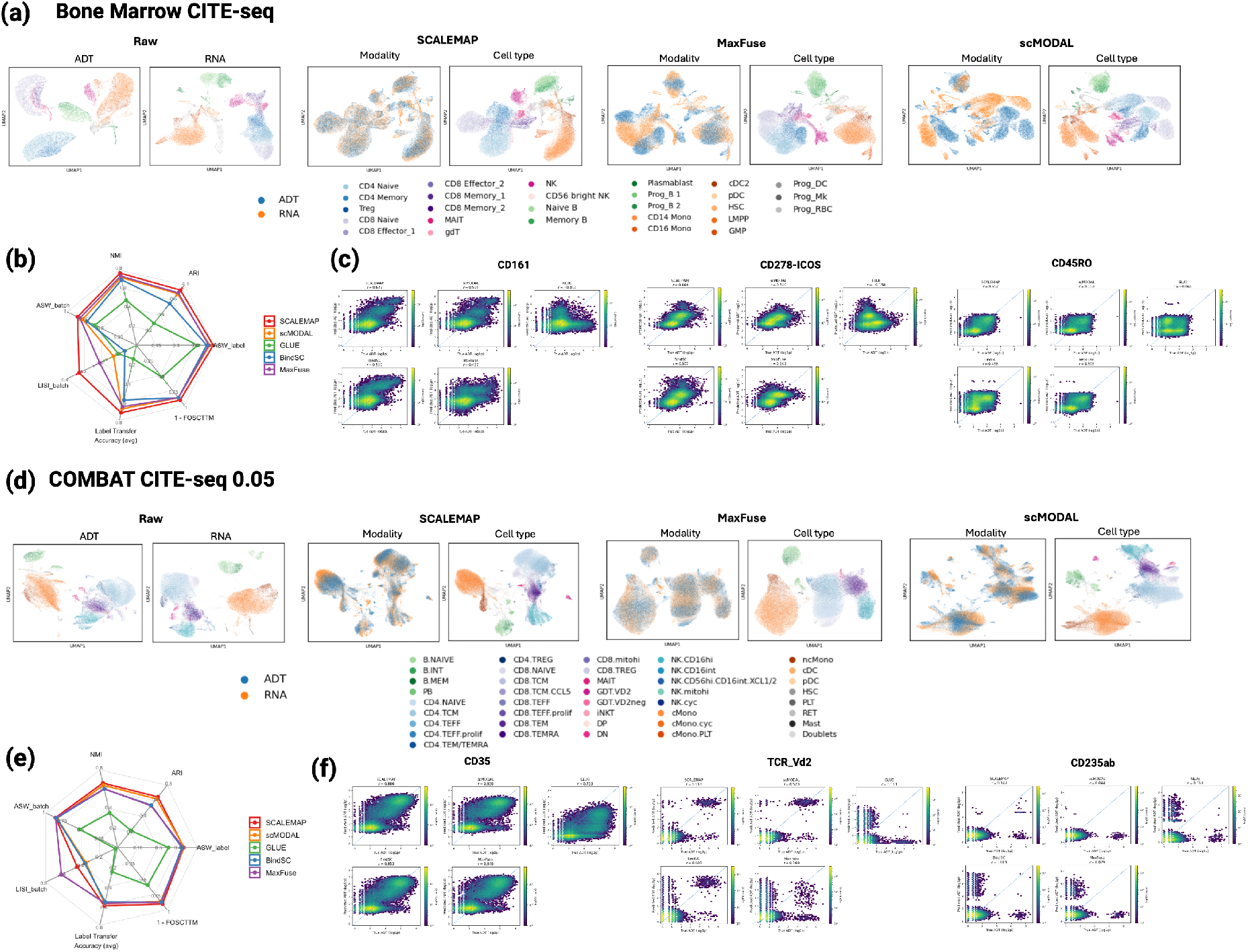
Integration quality and protein prediction under limited shared features and large-scale settings. (a) UMAP visualization of the Bone Marrow CITE-seq dataset. Raw RNA and ADT embeddings are shown alongside integrated results from SCALEMAP and MaxFuse. Points are colored by modality (left) and cell type (right). (b) Radar plot summarizing integration metrics on the Bone Marrow dataset. Each axis represents a metric, including biological conservation, batch correction, and modality alignment. (c) Marker-level RNA-to-protein prediction performance for selected Bone Marrow CITE-seq markers (CD161, CD278-ICOS and CD45RO), shown as predicted versus observed ADT abundance across cells and summarized by Pearson correlation. (d) UMAP visualization of the COMBAT CITE-seq (0.05 subset) dataset. (e) Radar plot of integration metrics on the COMBAT subset. (f) Marker-level RNA-to-protein prediction performance for selected COMBAT markers (CD35, TCR V*δ*2 and CD235ab), shown as predicted versus observed ADT abundance across cells and summarized by Pearson correlation.

We then examined whether numerical differences in RNA-to-protein prediction accuracy were reflected in visually discernible patterns in Bone Marrow CITE-seq. We focused on CD161, CD278-ICOS and CD45RO, selected to illustrate large between-method differences (Fig. 4c). SCALEMAP achieved the highest Pearson correlation for all three markers. The density-colored predicted-versus-observed scat-terplots visually illustrated closer agreement for SCALEMAP than scMODAL for CD278-ICOS, closer agreement for SCALEMAP than for MaxFuse for CD161 and CD278-ICOS, closer agreement than for BindSC for CD161 and CD45RO. Overall, SCALEMAP and scMODAL produced broadly more similar prediction patterns for the selected proteins, consistent with the earlier per-protein correlation comparisons (Fig. 3e). Additional visual difference can be observed when stratified by cell types, as illustrated by CD278-ICOS prediction in the CD8 Effector population (Fig. S4a-b).

In the subsetted COMBAT dataset, SCALEMAP produced an embedding with good RNA–ADT mixing and preservation of major immune cell populations, whereas MaxFuse and scMODAL showed different trade-offs between modality mixing and refined cell subpopulation organization (Fig. 4d). scMODAL performed the second best, with comparable biological conservation and modality alignment to SCALEMAP, which were 2.5% and 4.6% lower respectively, but showed weaker modality mixing, with a 21.4% lower LISI batch score than SCALEMAP. MaxFuse achieved the best modality mixing, with a 39.4% higher LISI batch score than SCALEMAP, but SCALEMAP achieved a 20.8% higher ARI than MaxFuse (Fig. 4e). Using the same focused local cross-modality kNN diagnostics, SCALEMAP better aligned refined T/NK and monocyte/DC populations (Fig. S3).

We also examined three representative COMBAT protein markers (Fig. 4f). CD235ab illustrated large differences between SCALEMAP and BindSC or scMODAL, while TCR V*δ*2 illustrated differences relative to MaxFuse. CD35 was included as a marker predicted with high correlation by all methods. SCALEMAP achieved the highest or second-highest Pearson correlation for all three markers. For CD35, the upper density region appeared more diagonally elongated for SCALEMAP and BindSC, whereas scMODAL showed a flatter pattern. For TCR V*δ*2, high predictions at low observed abundance were visually less prominent for SCALEMAP and scMODAL than for MaxFuse. For CD235ab, the upper-left region containing high predictions at low observed abundance was more abundant for BindSC and MaxFuse than for SCALEMAP and scMODAL. SCALEMAP also showed a small percent of cells with jointly high observed and predicted CD235ab abundance, although its overall correlation remained low. As in Bone Marrow CITE-seq, cell-type-specific views also showed more differences between SCALEMAP and scMODAL. For example, within 126 pDC cells, the fitted CD235ab trends had opposite signs, with Pearson correlations of 0.136 for SCALEMAP and −0.078 for scMODAL (Fig. S4c-d).

Full benchmark visualizations and metric summaries for the Bone marrow dataset, COMBAT subset are provided in Fig. S5. Fig. S6. Detailed visualizations for other datasets are also provided in Fig. S7-Fig. S10.

### 2.5 SCALEMAP achieves robust performance for RNA–ATAC integration across diverse TEA-seq datasets

We next evaluated RNA–ATAC integration using three benchmark datasets: PBMC TEA-seq, PBMC ASAPseqCtrl and PBMC ASAPseqStim (Supplementary Section 1). Compared with the RNA–protein benchmarks, the relative ranking of methods changed, with BindSC ranking second and SCALEMAP retaining the highest aggregate overall score (Fig. 5a). Across the three datasets, SCALEMAP also ranked first in aggregate biological conservation and batch correction, and second in modality alignment behind BindSC. SCALEMAP combined runtimes of a few minutes with peak memory use below 0.3 GiB. scMODAL required substantially longer training times, whereas MaxFuse and BindSC used more memory (Fig. 5b). UMAP comparisons with BindSC showed more uniform RNA–ATAC co-localization within shared cell populations under SCALEMAP (Fig. 5c–e).

**Fig. 5.**
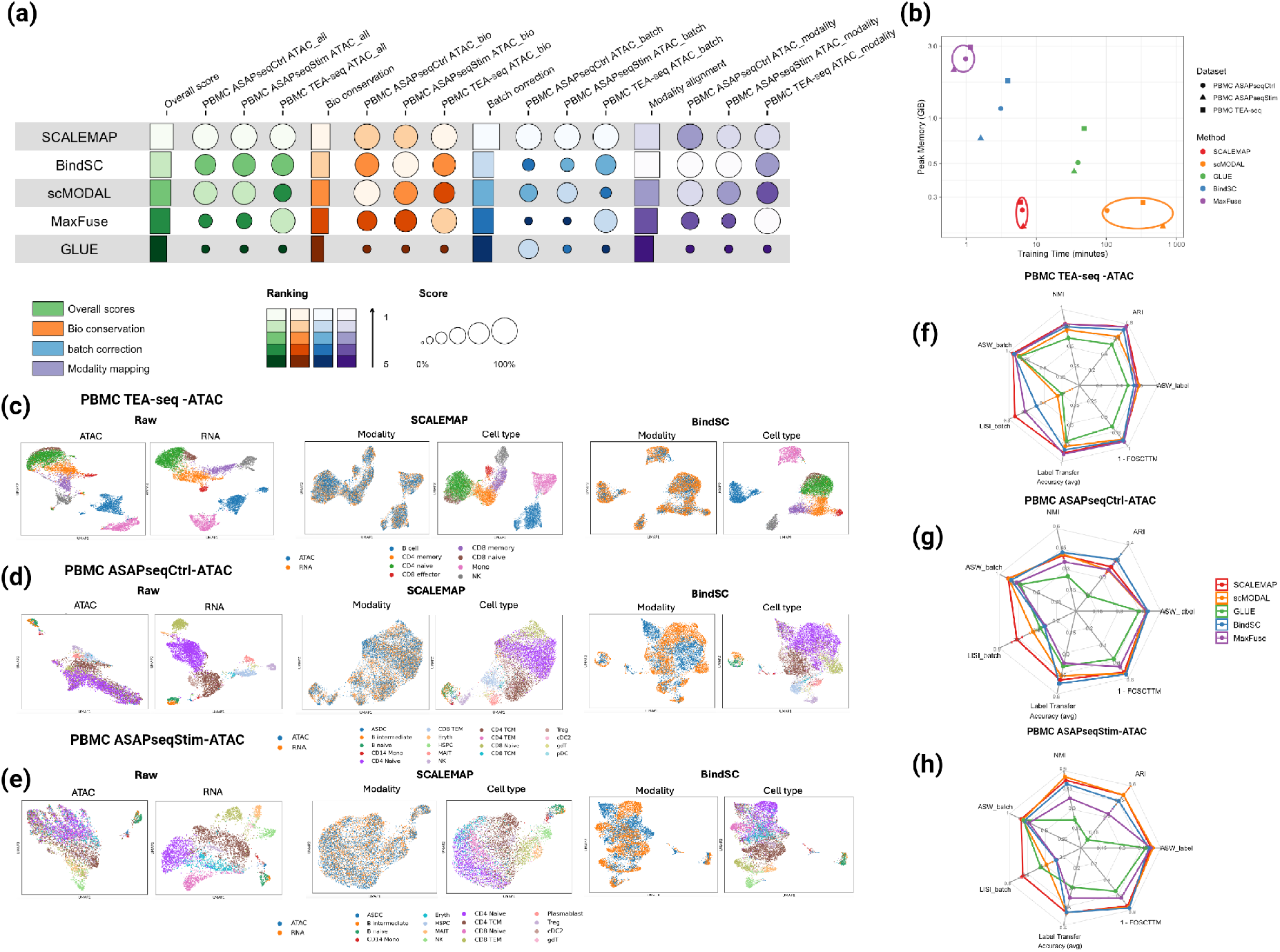
RNA–ATAC integration performance across TEA-seq datasets. (a) Bubble plot summarizing integration performance across three RNA–ATAC datasets. Circle size represents the magnitude of the score (linearly scaled), and color indicates metric category: green for overall score, orange for biological conservation, blue for batch correction, and purple for modality alignment. Methods are ranked based on their overall score. (b) Runtime and peak memory comparison across methods and RNA–ATAC datasets. (c–e) UMAP visualizations of raw and integrated embeddings for PBMC TEA-seq (c), PBMC ASAPseqCtrl (d), and PBMC ASAPseqStim (e). For each dataset, SCALEMAP is shown together with the best-performing non-SCALEMAP method by overall score to provide a focused visual comparison. Points are colored by modality (RNA vs ATAC) and cell type. (f–h) Radar plots summarizing integration metrics for PBMC TEA-seq (f), PBMC ASAPseqCtrl (g), and PBMC ASAPseqStim (h). Each axis represents a metric, including biological conservation, batch correction, and modality alignment, and each line corresponds to a method.

The relative performance of scMODAL and MaxFuse varied across datasets. scMODAL was more competitive on the ASAP-seq benchmarks than on PBMC TEA-seq. On PBMC TEA-seq, SCALEMAP achieved higher overall and category-level scores than scMODAL (Fig. 5f). The scMODAL UMAP showed more modality-stratified regions and less distinct separation between CD8 naive and CD4 naive cells (Supplementary Fig. S11). MaxFuse ranked second on PBMC TEA-seq but performed less well on the two ASAP-seq benchmarks (Fig. 5a). On PBMC ASAPseqCtrl, SCALEMAP achieved higher overall and batch-correction scores than MaxFuse (Fig. 5g). The MaxFuse embedding showed more modality-stratified regions and less distinct organization of T-cell subpopulations (Supplementary Fig. S12). In PBMC ASAPseqStim, annotated cell types were less clearly separated for the ATAC-seq modality(Fig. 5e). After integration, NMI values remained modest across methods, with the highest values around 0.4 (Fig. 5h). Despite this, SCALEMAP remains a top-tier method across all metrics (Fig. 5h).

### 2.6 Self-mapping is robust to small batch size

In methods that construct mutual nearest-neighbor (MNN) anchors within mini-batches or its variants, larger batches expand the candidate pool for potentially better cross-modality matching. This may increase the chance of sampling biologically corresponding cells, but also raises more computational and memory requirements. Self-mapping provides a cross-modality target derived from the same input cell for alignment through direct and learned cross-modality translation, which helps us to avoid searches for cross-modality cell matches in the observed data. Intuitively, this reduces the need of large batch size as it does not requires the existence of corresponding cell from the other modality. To assess whether self-mapping offers a more robustness towards reduced sample size, we compared integration quality as we vary mini-batch sizes from 50 to 2,000 cells. The evaluation covered six RNA–protein and three RNA–ATAC benchmark tasks (Fig. 6). We selected scMODAL as a leading-performing comparator whose within-batch MNN construction contrasts directly with SCALEMAP’s self-mapping strategy. Both methods use adversarial learning to encourage batch correction or modality mixing.

**Fig. 6.**
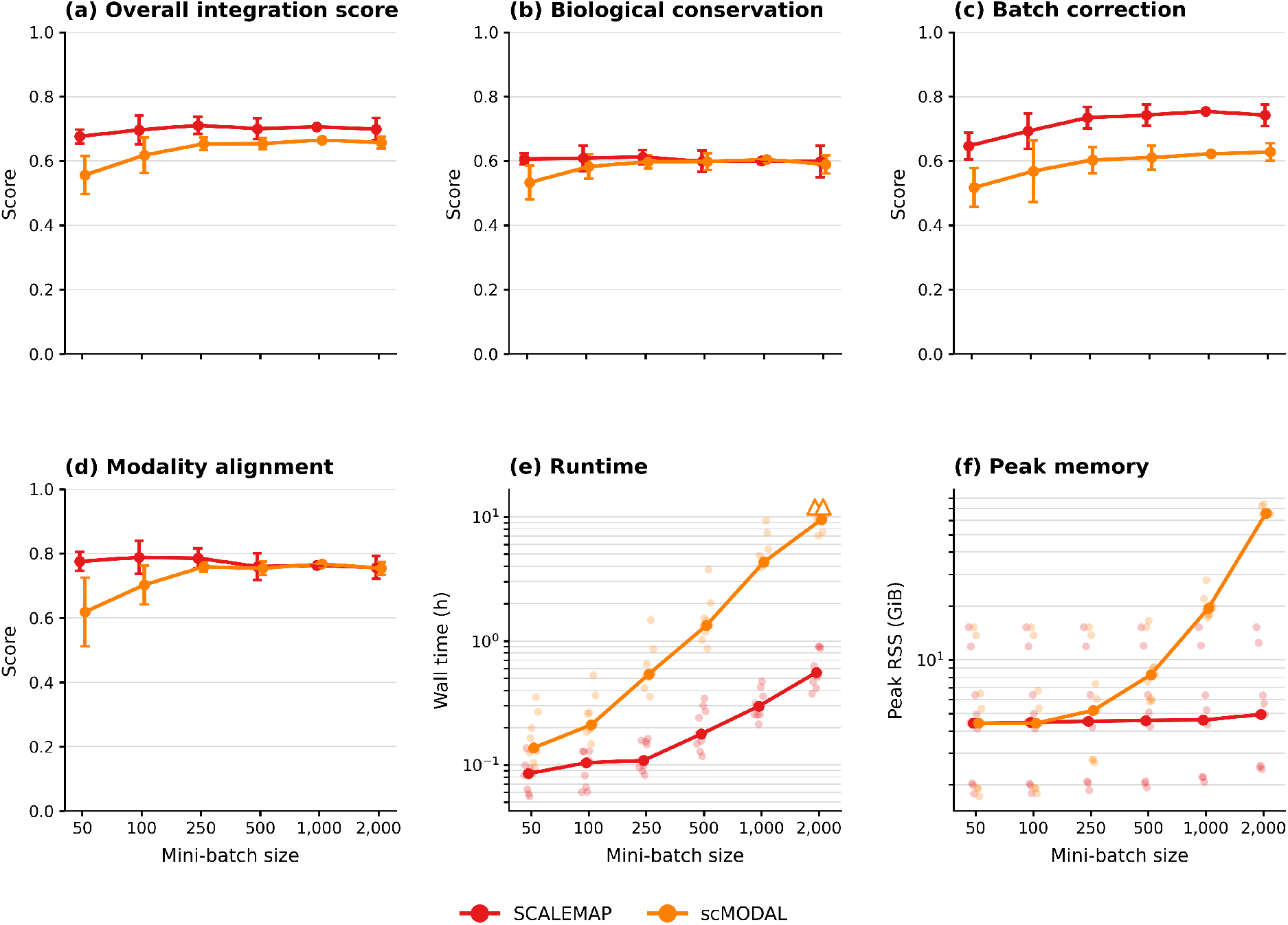
Mini-batch sensitivity and computational cost with varying mini-batch sizes across six RNA–ADT and three RNA– ATAC tasks. SCALEMAP and scMODAL were evaluated at mini-batch sizes from 50 to 2,000 cells. (a–d) Overall integration score (a), biological conservation (b), batch correction (c) and modality alignment (d). Solid lines show mean absolute scores across available tasks. Error bars are centered on these means and have half-width equal to the sample standard deviation across tasks of each task’s signed score change relative to the same method at batch size 1,000; they summarize variation in batch-size responses and are zero at the reference batch size. (e,f) Full-command wall time and peak resident memory, shown on logarithmic axes. Faint points represent individual completed tasks and solid lines connect the medians. Open triangles in e mark the two scMODAL tasks that exceeded the 12-h limit at batch size 2,000. All nine SCALEMAP tasks completed at every batch size; scMODAL summaries at batch size 2,000 include the seven completed tasks. Individual task trajectories are shown in Supplementary Fig. S14.

Across the nine benchmark tasks, SCALEMAP was less sensitive to reduced mini-batch size than scMODAL. As scMODAL did not complete training for two tasks at 2000 minibatch size, we used each tools’ respective performance at batch size 1,000 as the common reference to evaluate their potential performance decline associated with small batch size. SCALEMAP retained 95.8% of its aggregate overall score at batch size 50, compared with 83.6% for scMODAL. The difference in retention was greater for biological conservation and modality alignment than for batch correction. SCALEMAP retained 101.1% of its aggregate biological-conservation score and 101.0% of its modality-alignment score, compared with 88.1% and 73.2% for scMODAL, respectively. Aggregate batch-correction retention was closer between the methods, at 85.7% for SCALEMAP and 83.1% for scMODAL. Using task-specific reference scores, SCALEMAP showed higher retention than scMODAL in eight of nine biological-conservation comparisons and all nine modality-alignment comparisons. For batch correction, SCALEMAP showed higher retention in four of nine tasks (Fig. S14).

This performance differences originated from SCALEMAP’s robustness in bio-conservation and modality alignment: corresponding aggregate retention values were 101.1% versus 88.1% for biological conservation, and 101.0% versus 73.2% for modality alignment. The batch mixing trend were more similar between SCALEMAP and scMODAL: the corresponding retention values were 85.7% versus 83.1% for batch correction. At the task level, SCALEMAP had higher batch-size-50 retention in eight of nine biological-conservation and all nine modality-alignment comparisons, and four of nine batch-correction and(Fig. S14).

Computationally, all SCALEMAP runs completed within 12 h, whereas scMODAL exceeded this limit on RNA-ADT and RNA-ATACseq tasks on PBMC TEA-seq dataset at batch size 2,000 (Fig. 6a-b and Fig. S14). Among the seven tasks completed by both methods at this batch size, mean runtimes were 9.39 h for scMODAL and 0.60 h for SCALEMAP. Mean peak memory use was 67.75 GiB and 6.88 GiB, respectively. Runtime increased with batch size for both methods approximately linearly as batch size became large, but their memory-use trajectories differed. scMODAL’s peak memory rose sharply at larger batch sizes, whereas SCALEMAP’s remained comparatively stable up to a batch size of 2,000.

## 3 Discussion

We introduce SCALEMAP, a model for efficient diagonal integration through self-mapping. Biological feature links help to constrain each cell’s cross-modal translation. In matching-based approaches, feature links support the discovery of relationships between observed cells [23–25]. In contrast, self-mapping uses them to supervise translation directly and to give every sampled cell its own positive alignment target, without first identifying a partner in the other modality. This creates cell-specific alignment target without requiring paired measurements or inferred partners among available observed cells from the other modality.

Across six RNA-ADT integration datasets, SCALEMAP achieved the highest aggregate overall integration score and competitive embedding-based protein transfer (Fig. 3). In bone marrow and COMBAT where only very limited linkage informatin is available, integrated embedding or representation space preserved distinctions among selected refined immune and progenitor populations while bringing the modalities together (Fig. 4). SCALEMAP showed a clear advantage computationally when obtaining these integrated representations.

The RNA-ATAC experiments extend this evaluation to a second modality pairing (Fig. 5). SCALEMAP retained the highest aggregate overall score across these tasks, although the strongest alternative varied across metrics. Together with the RNA-ADT results, this supports the applicability of self-mapping across different types and densities of biological feature correspondence.

Self-mapping supplies biologically directed alignment constraints that complement adversarial modality mixing/batch correction used in SCALEMAP and many other prior tolls including scMODAL[25]. In the batch size sensitivity experiments, SCALEMAP preserved biological conservation and modality alignment more consistently than scMODAL, whereas modality mixing showed more similar sensitivity between the methods (Fig. 6). This pattern is consistent with a cell-specific alignment target that remains available as batch size decreases.

One limitation of self-mapping is that it still relies on predefined feature linkage to constrain across modality mapping, although through a robust contrastive learning. SCALEMAP currently defines its self-mapping targets using a fixed set of linked features. Earlier methods improve cross-modal correspondence through iterative cell matching or joint cell-and-feature alignment[24]. These approaches motivate extending self-mapping to learn link-specific weights or refine the feature comparison map under biological constraints. Such an extension could accommodate variation in link informativeness across cell states while retaining a target defined for each input cell.

In summary, SCALEMAP demonstrates that effective diagonal integration through self-mapping as a new modality alignment mechanism in horizontal integration. By replacing anchor search with cell-specific self-mapping, it provides a new way that balances integration quality, cross-modality prediction and computational efficiency.

## 4 Methods

### 4.1 Design and optimization of SCALEMAP

SCALEMAP takes two single-cell modalities as input, represented by 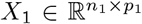 and 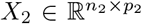, where the two modalities may contain different numbers of cells and different feature spaces.Each modality has a modality-specific input projection, followed by a shared encoder trunk that maps cells into a common latent space. Each modality also has its own decoder, allowing the model to reconstruct the observed modality and translate each cell into the other modality. Specifically, let modality *m* ∈ {1, 2} denote the two input modalities. For an input cell 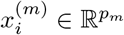, the encoder is defined as a modality-specific input layer followed by a shared encoder trunk:

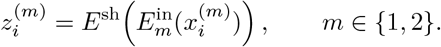

The modality-specific input mapping

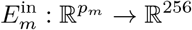

is implemented as a Linear*→*ReLU*→*BatchNorm block. The shared encoder trunk

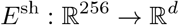

consists of

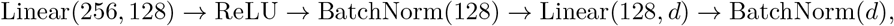

and is applied identically to both modalities. When samples from both modalities are used in the same forward pass, their modality-specific hidden representations are concatenated and jointly normalized through the shared trunk before being separated back into modality-specific latent representations.

Each modality has a decoder

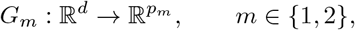

implemented as

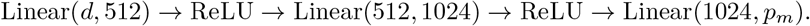

For a latent code 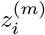, decoder *G*_*m*_ reconstructs the observed modality,

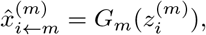

while decoder *G*_*¬m*_ generates the cross-modality prediction,

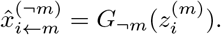

This architecture enables bidirectional cross-modality translation while learning a shared latent representation for both modalities.

To provide biologically meaningful correspondence between modalities, we use a curated correspondence table containing *s* linked feature pairs. This table defines shared-feature matrices 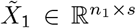 and 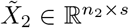. For RNA–protein integration, the linked features pair measured surface proteins with the expression of their corresponding coding genes. For RNA–ATAC integration, the linked features pair gene expression with gene activity scores derived from chromatin accessibility. The training objective is organized into three components: robust cell representation learning, modality mixing, and self-mapping through shared-feature contrastive learning. The first component ensures that the encoder–decoder path-ways learn stable and informative cell representations; the second component reduces modality-driven separation in the latent space; and the third component provides the main cross-modality mapping signal by matching each translated profile to the linked-feature representation of its own input cell.

#### Robust Cell Representation Learning

The first objective trains the encoder–decoder network to learn robust cell representations. It combines within-modality reconstruction, latent autoencoding consistency after cross-modality translation, and an augmented-view contrastive regularization term. These terms are grouped as one representation-learning component because they all support robust learning of the latent cell representation rather than directly enforcing cross-modality matching.

For each cell, SCALEMAP reconstructs the observed modality from its latent representation:

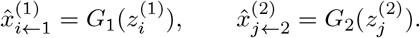

The reconstruction loss is

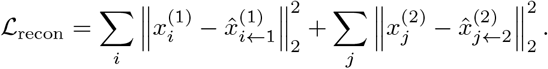

To ensure that cross-modality decoding preserves the latent identity of the input cell, SCALEMAP reencodes each translated profile and penalizes deviations from the original latent representation. For a cell 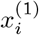, the model first translates it into modality 2:

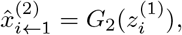

and then re-encodes the translated profile:

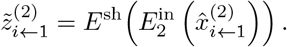

The same procedure is applied in the opposite direction:

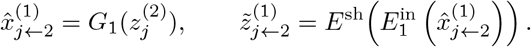

The latent consistency loss is

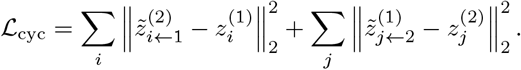

This term follows the same principle as cross-domain autoencoding consistency used in prior deep multiomics integration frameworks [25], but in SCALEMAP it is grouped with the representation-learning objective rather than treated as a separate alignment component.

To further improve robustness to sequencing-depth variation and local perturbations, SCALEMAP also uses augmented views of each cell. This follows the general practice of contrastive representation learning, where stochastic augmentations are used to encourage stable embeddings under realistic input perturbations [29–31]. For each modality *m*, two independently perturbed views of the same input cell are

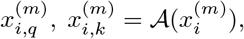

where *A*(·) denotes a stochastic sequencing-depth perturbation operator. In practice, this perturbation is implemented by random binomial thinning or Poisson sampling. The two views are encoded as

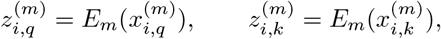

where *E*_*m*_ denotes the modality-specific input layer followed by the shared encoder.

The positive pair is formed by the two perturbed views of the same cell. To reduce false negatives among locally similar cells, SCALEMAP excludes cells from the same pre-integration neighborhood from the negative set. Let 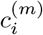 denote the pre-integration cluster assignment of cell *i* in modality *m*. The admissible negative set is

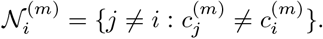

Using cosine similarity

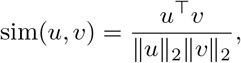

the augmented-view contrastive regularization for modality *m* is

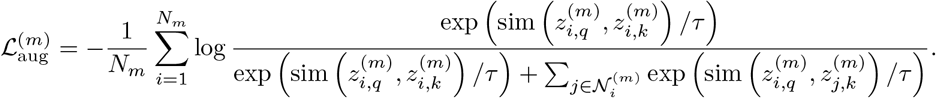

The regularization is averaged across modalities:

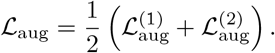

The full robust representation-learning objective is

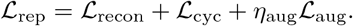

#### Modality Mixing via Adversarial Training

The second objective encourages modality mixing in the shared latent space. SCALEMAP uses a modality discriminator *D* that predicts the modality of each latent embedding, while the encoder is trained adversarially to make the embeddings less predictive of modality identity.

Let *v*_*i*_ be the one-hot modality label for cell *i*, and let *r*_*i*_ = *D*(*zi*) be the discriminator output. The discriminator minimizes the cross-entropy loss

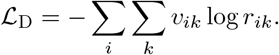

The discriminator is optimized to correctly distinguish modalities, whereas the encoder is optimized with the reversed objective to reduce modality-specific information in the latent space:

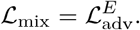

This adversarial component promotes modality mixing, but by itself it does not specify which biological states should correspond across modalities. Therefore, SCALEMAP combines it with the self-mapping objective described below.

#### Self-Mapping Through Shared-Feature Contrastive Learning

The third objective provides the main cross-modality mapping signal in SCALEMAP. Instead of using linked features to search for anchor cells across modalities, SCALEMAP uses the linked-feature correspondence to define a cell-specific target for its own translated profile. Let

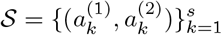

denote the ordered set of linked feature pairs. Let *Pm* extract the features from modality *m* corresponding to these links. For a cell 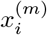, SCALEMAP extracts the linked-feature representation from its observed input:

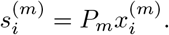

The same cell is translated into the other modality and then restricted to the corresponding linked features:

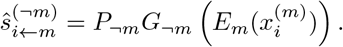

Both 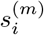 and 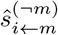 are represented in the same ordered linked-feature comparison space after modality-specific preprocessing. Therefore, the model can compare corresponding biological links without treating raw measurements from different assays as identical quantities.

For each modality, SCALEMAP uses an InfoNCE-style contrastive loss in which the translated linked-feature vector from cell *i* is encouraged to match the linked-feature representation of the same input cell more closely than translated vectors from other cells in the same minibatch:

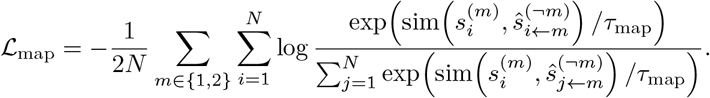

This self-mapping loss supplies diagonal supervision without requiring matched cells from the other modality. It is the main feature-correspondence-guided mapping component of SCALEMAP.

#### Model Optimization

SCALEMAP is trained by alternating between updates of the modality discriminator and updates of the encoder–decoder network. The final encoder–decoder objective contains three top-level components:

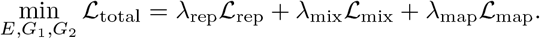

Here, ℒ_rep_ trains robust encoder–decoder pathways through reconstruction, autoencoding consistency and augmented-view regularization; ℒ_mix_ promotes modality mixing in the latent space through adversarial training; and ℒ_map_ performs self-mapping through shared-feature contrastive learning.

The discriminator *D* is trained separately by minimizing

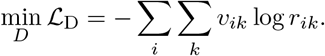

In each training iteration, balanced minibatches are sampled from the two modalities. The discriminator is first updated using the current latent embeddings. The encoder and decoders are then updated using the full objective above.

We fix the hyperparameter values as following,

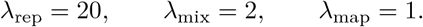

Within ℒ_rep_, the augmented-view regularization uses a fixed internal weight

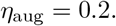

The encoder and decoders are optimized using Adam with learning rate 10^™3^, while the discriminator is optimiz
ed using Adam with learning rate 2 × 10^™3^. A weight decay of 10^™3^ is used for both optimizers. Training is performed for 2,000 iterations. At each iteration, minibatches are sampled separately from the two modalities, with 250 cells sampled from each modality. To stabilize adversarial optimization, the discriminator is updated seven times before each encoder–decoder update. Gradient clipping with maximum norm 5 is applied during the encoder–decoder update.

#### Model Outputs

After training, SCALEMAP outputs the integrated latent embeddings

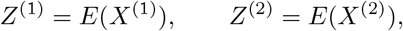

as well as bidirectional cross-modality predictions

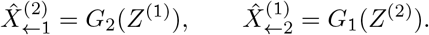

The integrated embeddings are used for visualization, clustering, cell-type annotation and benchmarking, while the cross-modality predictions provide translated feature profiles for downstream biological interpretation.

### 4.2 Protein Prediction

To ensure comparable RNA-to-protein prediction evaluation across methods, protein abundance was predicted from each method’s integrated embedding using the same nearest-neighbor procedure. For each RNA-profile cell, we identified its 10 nearest protein-profile neighbors in the integrated space and averaged their observed protein measurements.

Supplementary results also additionally utilized random forest for protein prediction. Random forest-based RNA-to-protein prediction was also constructed using each method’s precomputed integrated embeddings. For each protein, a random forest with 300 trees was trained to predict observed protein abundance from the training cells’ ADT embeddings and then applied to the test cells’ RNA embeddings.

### 4.3 Evaluation Metrics

Integration performance was evaluated using biological conservation, batch correction and modality alignment metrics, as detailed in Supplementary Section 3. Biological conservation was summarized by NMI, ARI and cell-type ASW; batch correction was summarized by batch ASW and batch LISI; and modality alignment was summarized by label-transfer accuracy and 1 ™ FOSCTTM. Category scores were computed by averaging the corresponding metrics, and the overall score was computed as the average of the three category scores.

Because the refined subsets in Fig. S3 are small and not always clearly separated into discrete clusters, we used separate macro-averaged local cross-modality kNN diagnostics in the existing integrated embedding rather than ARI or NMI for this focused analysis. These diagnostics evaluate local RNA– ADT alignment within selected rare or refined populations. Specifically, we computed opposite-modality label agreement and opposite-modality hit@k within the selected subsets, as defined in Supplementary Section 3. In Bone Marrow, SCALEMAP showed the strongest opposite-modality label agreement in both selected subsets. In COMBAT, SCALEMAP better aligned refined T/NK and monocyte/DC populations across modalities, with stronger opposite-modality label agreement and opposite-modality hit@k in the selected subsets. For the marker-level prediction analyses in Fig. 4c,f and all panels of Supplementary Fig. S4, we used Pearson correlation between predicted and observed marker abundance. The supplementary representative panels compute Pearson correlation within the displayed cell type.

Prediction accuracy for each protein was then measured by the Pearson correlation between the predicted and observed protein abundance across cells. Dataset-level protein prediction performance was summarized by the mean Pearson correlation across proteins.

## Acknowledgements

This work is supported by NSF DMS2310836, 5U01AI167892, and R01AI198335.

## Declarations

- Data availability: See Supplementary Material, Suppl Section 1
- Code availability: https://github.com/ch2343/SCALEMAP.git

