## Supplementary material for "Scalable Multi-omics Diagonal Integration from Self-Mapping": Supplemetary Note

Chuan He[1*] and Leying Guan[1,2*]

[1*]Department of Biostatistics, Yale School of Public Health, New Haven, 06511, CT, US.
[2]Program in Computational Biology and Bioinformatics, Yale University, New Haven, 06511, CT, US.

#### Contents

**1 Dataset information** **2**

**2 Training Algorithms** **4**

**3 Evaluation Metrics** **5**

**4 Mini-batch Sensitivity Benchmark Design and Measurements** **8**

**5 Supplementary Figures** **9**

### 1 Dataset information

We evaluated SCALEMAP on six publicly available single-cell multi-omics datasets spanning different tissues, technologies, and modality combinations. For each dataset, we extracted paired modalities (RNA–ADT or RNA–ATAC) and constructed corresponding shared feature spaces as described in the preprocessing section.

##### *PBMC CITE-seq*

This dataset consists of human peripheral blood mononuclear cells (PBMCs) collected from eight donors before and after COVID-19 vaccination. We used a subset following the Wang et al. [1] setup to reduce inter-individual batch effects. The dataset contains 18,034 cells with 20,729 RNA features and 228 protein (ADT) features. A total of 221 RNA–protein correspondence pairs were identified, forming the shared feature space.

##### *Bone Marrow CITE-seq*

This dataset was introduced in *Comprehensive Integration of Single-Cell Data* (Stuart et al., Cell 2019) and contains human bone marrow samples projected onto the Human Cell Atlas (HCA) reference. The subset used here includes 30,672 cells with 17,009 RNA features and 25 ADT features. A total of 25 RNA–protein correspondence pairs were constructed, representing a highly sparse shared feature setting.

##### *COMBAT CITE-seq*

The COMBAT dataset (GSE35216673) consists of large-scale PBMC samples collected from COVID-19 and sepsis cohorts. We use full data with 831,742 cells for scalable evaluation and a subset containing 41,807 cells with 2,057 RNA features and 192 ADT features for full metrics evaluation across methods. Due to differences in naming conventions, a dataset-specific correspondence table was constructed, resulting in 57 shared RNA–protein features.

##### *TEA-seq (RNA–ADT / RNA–ATAC).*

The TEA-seq dataset (GSE158013) provides trimodal measurements of RNA, surface proteins (ADT), and chromatin accessibility (ATAC) from the same cells. We extract paired RNA–ADT and RNA–ATAC subsets for evaluation. The dataset contains 7,437 cells with 18,352 RNA features, 46 ADT features, and 18,352 ATAC features. For RNA–ADT, 46 correspondence pairs were identified, while for RNA–ATAC, all features were matched via gene name intersection.

##### *D22 (ASAP-seq control).*

The D22 dataset is derived from GSE156478 and corresponds to control PBMC samples. It contains 6,408 cells with 12,408 RNA features and 210 ADT features. For RNA–ADT integration, 121 shared features were constructed. For RNA–ATAC integration, 10,007 shared features were obtained via feature-name matching between RNA and ATAC.

##### *D23 (ASAP-seq stimulated).*

The D23 dataset, also from GSE156478, corresponds to stimulated PBMC samples. It includes 4,606 cells with 12,773 RNA features and 210 ADT features. A total of 126 RNA–ADT shared features were identified. For RNA–ATAC integration, 10,323 shared features were obtained via name intersection.

#### Dataset Summary

**Table S1** Summary of datasets and feature statistics used in this study.

| Dataset | Modality | Cells | RNA features | Mod2 features | Shared | GSE |
|---|---|---|---|---|---|---|
| Bone Marrow CITE-seq | RNA–ADT | 30,672 | 17,009 | 25 | 25 | GSE128639 |
| PBMC ASAPseqCtrl | RNA–ADT | 6,408 | 12,408 | 210 | 121 | GSE156478 |
| PBMC ASAPseqStim | RNA–ADT | 4,606 | 12,773 | 210 | 126 | GSE156478 |
| PBMC CITE-seq | RNA–ADT | 18,034 | 20,729 | 228 | 221 | GSE164378 |
| COMBAT CITE-seq | RNA–ADT | 41,807 | 2,057 | 192 | 57 | GSE35216673 |
| PBMC TEA-seq | RNA–ADT | 7,437 | 18,352 | 46 | 46 | GSE158013 |
| PBMC ASAPseqCtrl | RNA–ATAC | 6,408 | 12,408 | 20,149 | 10,007 | GSE156478 |
| PBMC ASAPseqStim | RNA–ATAC | 4,606 | 12,773 | 20,599 | 10,323 | GSE156478 |
| PBMC TEA-seq | RNA–ATAC | 7,437 | 18,352 | 18,352 | 18,352 | GSE158013 |

### 2 Training Algorithms

***Notation.***

$E$ denotes the shared encoder, $G$ the decoder(s), and $D$ a discriminator. $x_A, x_B, x_C$ are minibatches from modalities $A, B, C$ with cluster labels $\ell_A, \ell_B, \ell_C$. $\textsc{MakeDepthViews}(\cdot)$ produces two stochastic views. $\textsc{InfoNCE}(\cdot)$ excludes same-label negatives.

**Algorithm 1** Two-Modality Training (A/B)

**Require:** Data $(A, B)$, predictors Pred, steps $T$
**Require:** Weights $\lambda_{\text{AE}}, \lambda_{\text{RE}}, \lambda_{\text{GAN}}, \lambda_{\text{Align}}, \lambda_{\text{noise}}$

```
Initialize encoder E, decoder G, discriminator D
for t = 1 to T do
    Sample minibatches (x_A, x_B) and labels (ℓ_A, ℓ_B)
    (x_A^q, x_A^k) ← MakeDepthViews(x_A)
    (x_B^q, x_B^k) ← MakeDepthViews(x_B)
    L_noise ← 1/2 [InfoNCE(E(x_A^q), E(x_A^k), ℓ_A) + InfoNCE(E(x_B^q), E(x_B^k), ℓ_B)]
    (z_A, z_B) ← E(x_A, x_B)
    (x̂_{B→A}, x̂_{A→B}), (x̂_A, x̂_B) ← G(z_B, z_A), G(z_A, z_B)
    L_AE ← ‖x̂_A − x_A‖^2 + ‖x̂_B − x_B‖^2
    L_RE ← ‖E(x̂_{B→A}) − z_B‖^2 + ‖E(x̂_{A→B}) − z_A‖^2
    L_Align ← 0.2 [Contrastive(Pred(x_A), Pred(x̂_{A→B})) + Contrastive(Pred(x_B), Pred(x̂_{B→A}))]
    Update D for 5 steps using D([z_A, z_B])
    L_GAN ← −𝔼[D([z_A, z_B])]
    L_G ← λ_GAN L_GAN + λ_AE (L_AE + L_RE) + λ_Align L_Align + λ_noise L_noise
    Update (E, G) using L_G
end for
```

### 3 Evaluation Metrics

#### Overview of Basic Integration Evaluation Metrics

We evaluate integration performance using three complementary aspects: **biological conservation**, **batch correction**, and **modality alignment**. Biological conservation assesses the preservation of cell-type structure, batch correction measures the removal of technical variation across modalities or batches, and modality alignment evaluates the consistency of corresponding cells across modalities. In total, we consider seven metrics: three for biological conservation (NMI, ARI, ASW_label), two for batch correction (ASW_batch, LISI_batch), and two for modality alignment (label transfer accuracy and FOSCTTM).

**Table S2** Summary of evaluation metrics and score aggregation.

| Metric | Description |
|---|---|
| **Biological Conservation** | |
| Normalized Mutual Information (NMI) | Agreement between cell-type labels and clustering results. |
| Adjusted Rand Index (ARI) | Clustering accuracy adjusted for chance. |
| ASW_label | Measures cluster separation and cohesion based on cell types. |
| **Batch Correction** | |
| ASW_batch | Evaluates mixing of batches in the embedding space. |
| LISI_batch | Measures batch diversity in local neighborhoods. |
| **Modality Alignment** | |
| Label Transfer Accuracy | Accuracy of transferring labels across modalities (averaged bidirectionally). |
| FOSCTTM | Fraction of samples closer than the true match (reported as $1 - \text{FOSCTTM}$). |
| **Score Aggregation** | |
| Biological Conservation Score | Average of NMI, ARI, and ASW_label. |
| Batch Correction Score | Average of ASW_batch and LISI_batch. |
| Modality Alignment Score | Average of label transfer accuracy and ($1 - \text{FOSCTTM}$). |
| Overall Score | Average of biological conservation, batch correction, and modality alignment scores. |

#### Details on Biological Conservation Metrics

For cluster-based biological conservation metrics (NMI and ARI), clustering of the integrated data is required. Following [2], we apply Louvain clustering and select the resolution that maximizes the NMI score.

**Normalized Mutual Information (NMI).** NMI measures the agreement between predicted clusters and ground-truth cell-type labels. It is defined as:

$$\text{NMI}(U, V) = \frac{2I(U, V)}{H(U) + H(V)}$$

where $I(U, V)$ is the mutual information and $H(\cdot)$ denotes entropy. The score ranges from 0 (no agreement) to 1 (perfect agreement).

**Adjusted Rand Index (ARI).** ARI evaluates clustering similarity while correcting for random assignments [3, 4]. It ranges from 0 (random labeling) to 1 (perfect match).

**Cell Type Average Silhouette Width (ASW_label).** ASW measures cluster cohesion and separation:

$$ASW = \frac{1}{N} \sum_{i=1}^{N} \frac{b(i) - a(i)}{\max\{a(i), b(i)\}}$$

where $a(i)$ is the intra-cluster distance and $b(i)$ is the nearest-cluster distance. The final score is scaled to $[0, 1]$ via $(ASW + 1)/2$.

#### Details on Batch Correction Metrics

**ASW_batch.** ASW_batch evaluates how well cells from different batches are mixed:

$$ASW_{\text{batch}} = \frac{1}{N} \sum_{i=1}^{N} \frac{b(i) - a(i)}{\max\{a(i), b(i)\}}$$

where $a(i)$ measures within-batch distance and $b(i)$ measures distance to other batches. The final score is transformed as:

$$ASW_{\text{batch}}^{\text{final}} = 1 - |ASW_{\text{batch}}|$$

so that higher values indicate better mixing.

**LISI_batch.** LISI measures batch diversity within local neighborhoods using an inverse Simpson index [5]:

$$LISI = \frac{1}{\sum_{b=1}^{B} p(b)^2}$$

where $p(b)$ is the probability of batch $b$ within a neighborhood. Scores are scaled to reflect effective batch mixing.

#### Details on Modality Alignment Metrics

**Label Transfer Accuracy.** Label transfer accuracy evaluates how well cell-type labels can be transferred across modalities. For each modality, we perform nearest-neighbor classification in the integrated embedding and compute accuracy in both directions (e.g., RNA→ADT and ADT→RNA). The final score is the average of the two directions:

$$\text{Accuracy} = \frac{1}{2}(\text{acc}_{1\to 2} + \text{acc}_{2\to 1})$$

**FOSCTTM (Fraction of Samples Closer Than the True Match).** FOSCTTM measures how well corresponding cells are aligned across modalities [6]. For each cell, it computes the fraction of cells in the opposite modality that are closer than its true match:

$$FOSCTTM = \frac{1}{N} \sum_{i=1}^{N} \frac{\#\{j : d(i,j) < d(i,i^*)\}}{N}$$

where $i^*$ denotes the true paired cell. Lower values indicate better alignment, and we report 1−FOSCTTM so that higher scores correspond to better performance.

#### Protein Prediction and Cell-type-level Prediction Metrics

For RNA-to-protein prediction evaluation, all methods were compared using the same nearest-neighbor imputation procedure applied to their integrated embeddings. For each RNA-profile cell $i$, we identified the $k = 10$ nearest protein-profile cells $N_{10}(i)$ in the integrated space and predicted protein $g$ by averaging the observed protein abundance among these neighbors:

$$\hat{y}_{ig} = \frac{1}{|N_{10}(i)|} \sum_{j \in N_{10}(i)} y_{jg}.$$

For each protein, prediction accuracy was measured by the Pearson correlation between observed and predicted protein abundance across cells:

$$r_g = \frac{\sum_i (y_{ig} - \bar{y}_g)(\hat{y}_{ig} - \bar{\hat{y}}_g)}{\sqrt{\sum_i (y_{ig} - \bar{y}_g)^2} \sqrt{\sum_i (\hat{y}_{ig} - \bar{\hat{y}}_g)^2}}.$$

The dataset-level protein prediction score was the mean Pearson correlation across proteins.

Protein prediction within annotated cell types was evaluated by Pearson correlation (Supplementary Fig. S4). For method $m$, protein $g$, and cell type $c$, the within-cell-type correlation was

$$r_{cg}^{(m)} = \text{corr}\Big(\{y_{ig} : i \in c\}, \{\hat{y}_{ig}^{(m)} : i \in c\}\Big).$$

For the selected observed-versus-predicted examples, we additionally fitted an ordinary least-squares regression within each method, protein and cell type,

$$\hat{y}_{ig}^{(m)} = \alpha_{cg}^{(m)} + \beta_{cg}^{(m)} y_{ig} + \varepsilon_{ig},$$

where $\beta_{cg}^{(m)}$ is the fitted slope reported in the corresponding panel. Pearson $r$ measures the strength of linear association, whereas $\beta$ describes how strongly the predicted abundance changes with observed abundance. A positive slope indicates that higher observed values tend to receive higher predictions; a slope closer to one indicates better agreement in scale, although calibration also depends on the fitted intercept. The solid red line shows the fitted regression, and the dashed identity line represents exact agreement between observed and predicted abundance. Panels a and c show these correlations for CD278-ICOS and CD235ab, respectively, across the cell types displayed in Supplementary Fig. S3. The single-cell examples in panels b and d were chosen as the cell types with the largest between-method Pearson-correlation difference for the corresponding marker among these displayed cell types: CD8 Effector 1 for CD278-ICOS and pDC for CD235ab.

#### Focused Local Cross-Modality kNN Metrics for Refined Subsets

For the focused subcluster analysis in Fig. S3, we additionally computed two local cross-modality k-nearest-neighbor (kNN) diagnostics. These metrics were reported separately from the seven global benchmark metrics and the overall score aggregation. They were designed to evaluate local RNA–ADT alignment within rare or refined cell populations after subsetting cells from the already learned integrated embedding, without reclustering or recomputing the embedding.

Let $z_i$ denote the integrated embedding of cell $i$, $y_i$ its fine-grained cell-type label, and $m_i$ its modality label. For each selected subset, we identified the Euclidean kNN set $N_k(i)$ in the existing integrated space with $k = 30$, excluding cell $i$ itself. To directly evaluate cross-modality local alignment, we restricted the neighborhood to cells from the opposite modality:

$$O_k(i) = \{j \in N_k(i) : m_j \neq m_i\}.$$

The opposite-modality label agreement was defined as

$$A_i = \frac{1}{|O_k(i)|} \sum_{j \in O_k(i)} \mathbf{1}(y_j = y_i),$$

and the opposite-modality hit@k was defined as

$$H_i = \mathbf{1}\left(\exists j \in O_k(i) \text{ such that } y_j = y_i\right).$$

Cells without opposite-modality neighbors among their $k$ nearest neighbors were omitted from the corresponding opposite-modality average. Thus, $A_i$ measures the fraction of cross-modality neighbors with the same fine annotation, whereas $H_i$ is a more permissive retrieval-style metric that asks whether at least one matched cross-modality neighbor is present.

For each metric $Q_i \in \{A_i, H_i\}$, we first averaged over cells within each fine-grained cell type $c$, and then averaged equally over the $C$ selected cell types:

$$Q_{\text{macro}} = \frac{1}{C} \sum_{c=1}^{C} \frac{1}{n_c} \sum_{i:y_i=c} Q_i.$$

This macro-average prevents abundant cell types from dominating the focused evaluation and gives rare refined populations equal weight in the reported local metrics.

### 4 Mini-batch Sensitivity Benchmark Design and Measurements

We evaluated sensitivity to training mini-batch size across nine benchmark tasks: six RNA–ADT tasks (PBMC CITE-seq, Bone Marrow CITE-seq, COMBAT CITE-seq, PBMC TEA-seq, PBMC ASAPseqCtrl and PBMC ASAPseqStim) and three RNA–ATAC tasks (PBMC TEA-seq, PBMC ASAPseqCtrl and PBMC ASAPseqStim). SCALEMAP and scMODAL were each evaluated at batch sizes 50, 100, 250, 500, 1,000 and 2,000, giving 108 planned conditions. The integration seed was fixed at 10. Method-specific training durations were retained at 2,000 iterations for SCALEMAP and 5,000 iterations for scMODAL, and only the mini-batch size was changed within each method. Runtime was measured for the complete integration command, and peak memory was measured as maximum resident set size.

For each method, small-batch retention was evaluated relative to batch size 1,000, which was completed for all nine tasks by both methods. If $S_{m,c}(s)$ denotes the aggregate score for method $m$, score category $c$ and batch size $s$, retention was defined as

$$R^{\%}_{m,c}(s) = 100\frac{S_{m,c}(s)}{S_{m,c}(1000)}.$$

The same definition was applied to each task and score category, $R^{\%}_{d,m,c}(s) = 100 S_{d,m,c}(s)/S_{d,m,c}(1000)$, for the dataset-level comparisons reported in Results. This common reference separates sensitivity to batch-size reduction from differences in absolute performance between methods. In main-text Fig. 6a–d, solid lines show mean absolute scores across available tasks. To describe variation in batch-size responses separately from differences in task difficulty, error-bar half-widths were calculated using the same batch-size-1,000 reference as

$$E_{m,c}(s) = \mathrm{SD}_d \left[S_{d,m,c}(s) - S_{d,m,c}(1000)\right],$$

where $d$ indexes tasks, $m$ denotes the method and $c$ denotes the score category. The differences are signed score-point changes, and the sample standard deviation is calculated over tasks with both scores available. Error bars are centered on the mean absolute scores, summarize task-to-task variation in batch-size responses and are zero at batch size 1,000. Main-text Fig. 6e,f show individual completed tasks and median runtime and memory at each batch size. Supplementary Fig. S14 presents all task-level score and resource trajectories; its aggregate performance row retains the sample standard deviation of absolute scores across tasks. All nine SCALEMAP tasks completed at every tested batch size. All nine scMODAL tasks completed at batch sizes 50–1,000, whereas the PBMC TEA-seq RNA–ADT and RNA–ATAC tasks at batch size 2,000 exceeded the 12-h limit. The scMODAL batch-size-2,000 summaries therefore comprise the seven completed tasks.

### 5 Supplementary Figures

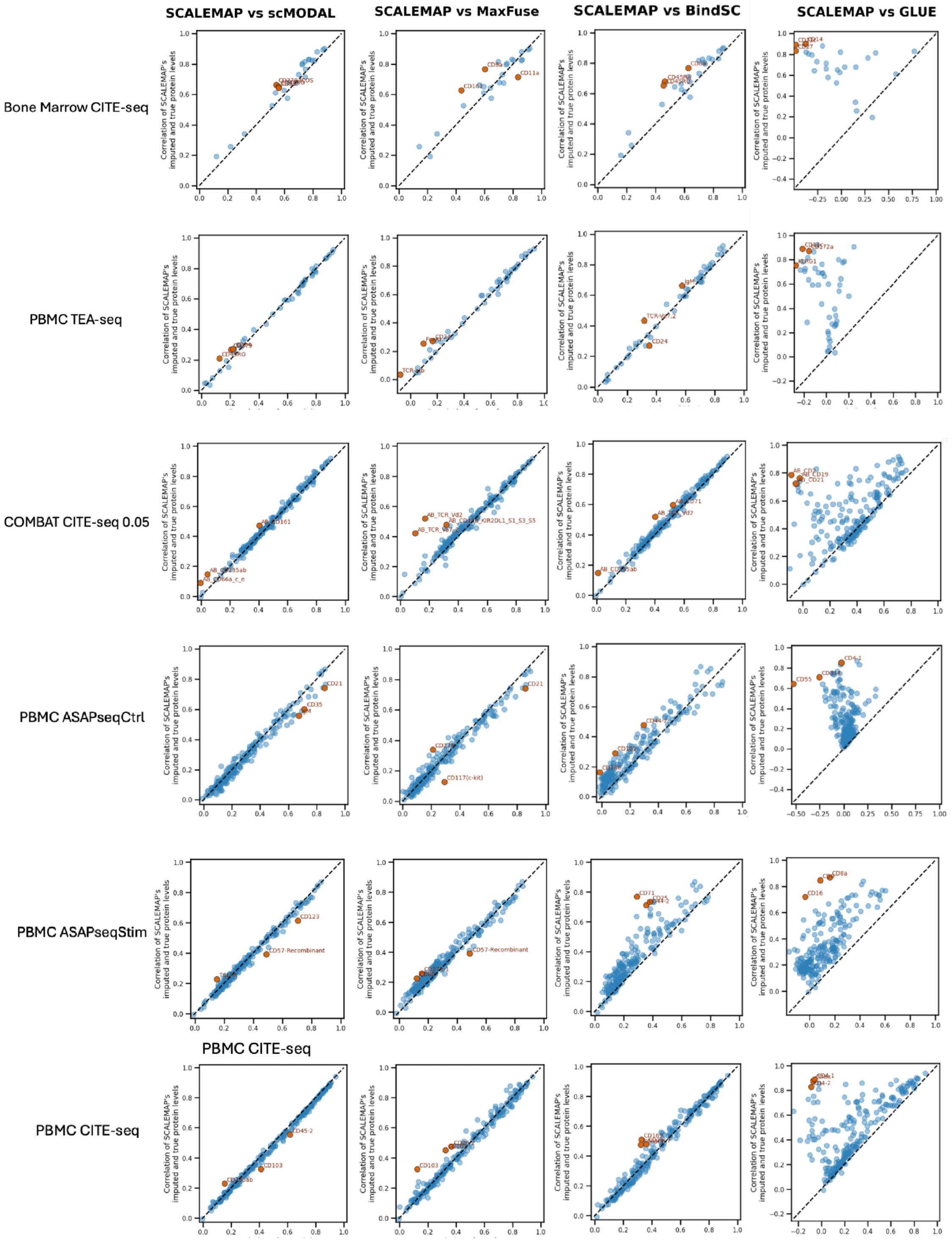


**Fig. S1** RNA-to-protein prediction using KNN-based imputation across six RNA–ADT datasets. Each point represents one protein, with its predicted-versus-observed Pearson correlation for SCALEMAP on the y-axis and for scMODAL, MaxFuse, BindSC or GLUE on the x-axis, as indicated above each column. Rows correspond to individual datasets. Dashed lines denote equal correlations; points above the line indicate higher correlation for SCALEMAP. Selected proteins with large between-method differences are labeled.

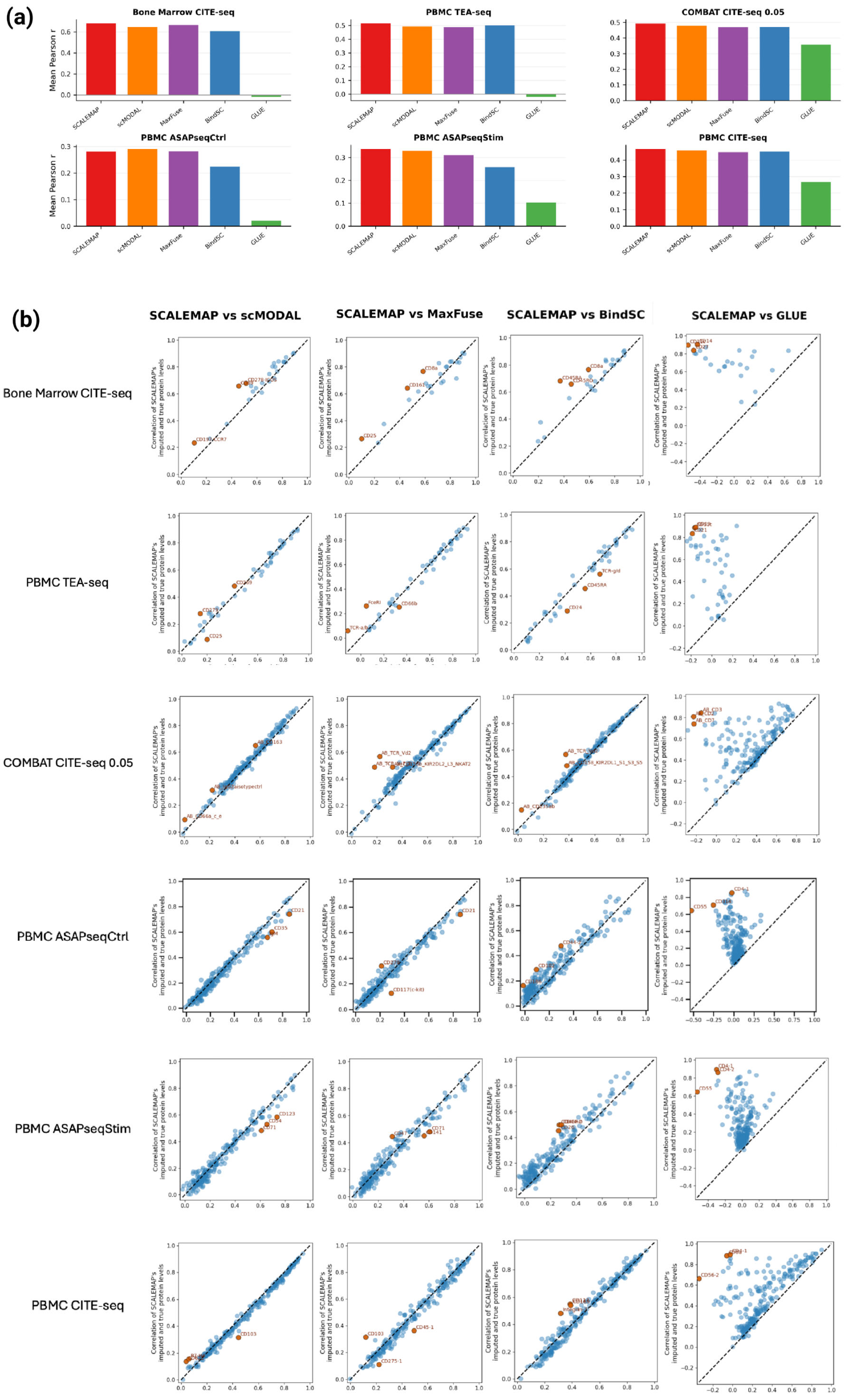


**Fig. S2** RNA-to-protein prediction using random forest-based imputation across six RNA–ADT datasets. (a) Mean predicted-versus-observed Pearson correlation across proteins for each integration method and dataset. (b) Protein-wise Pearson correlations for SCALEMAP (y-axis) versus scMODAL, MaxFuse, BindSC or GLUE (x-axis), as indicated above each column. Correlations were computed across test cells. Each point represents one protein, and rows correspond to individual datasets. Dashed lines denote equal correlations; selected proteins with large between-method differences are labeled.

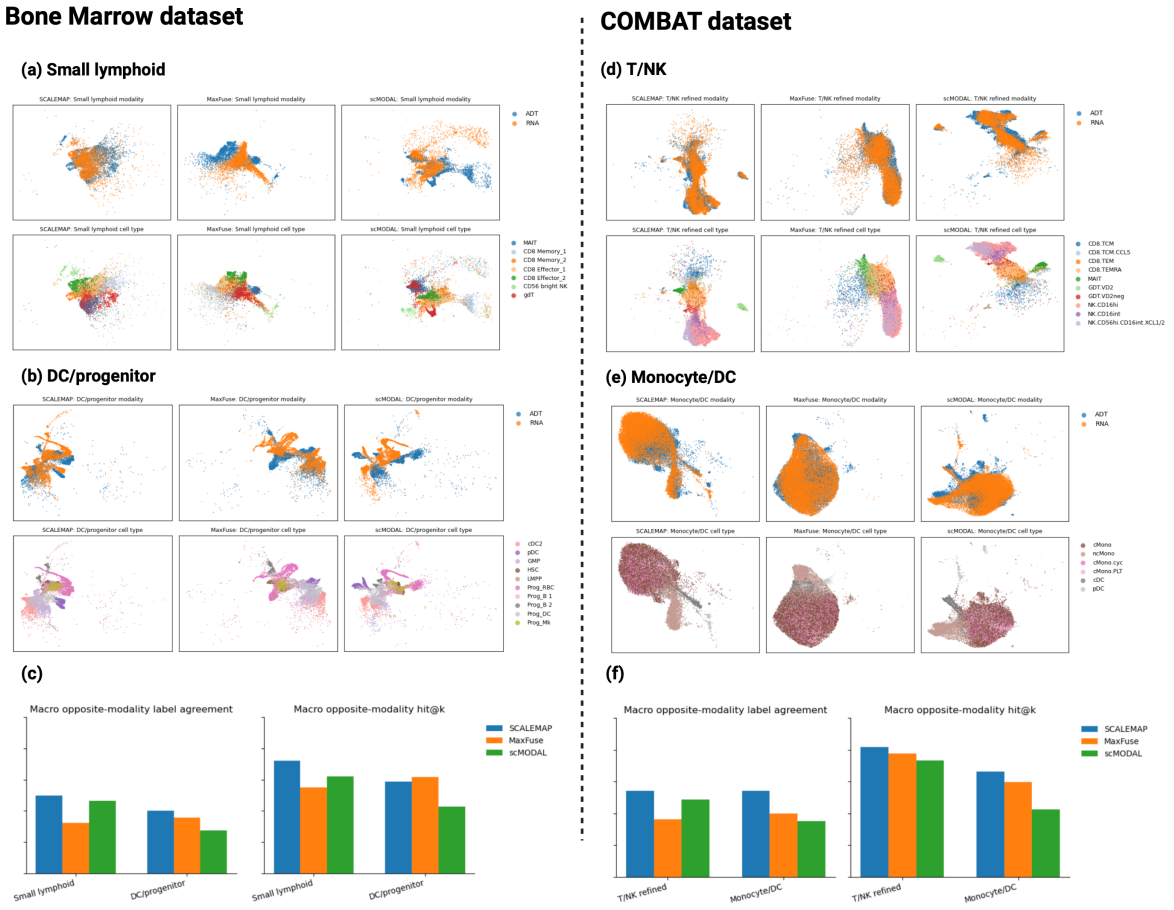


**Fig. S3** Focused subcluster analysis in the Bone Marrow and COMBAT datasets. (a,b) UMAP visualizations of selected small lymphoid and dendritic/progenitor-associated populations in the Bone Marrow CITE-seq dataset, colored by modality and cell type across SCALEMAP, MaxFuse and scMODAL. (c) Quantitative comparison of macro-averaged opposite-modality label agreement and opposite-modality hit@k for the selected Bone Marrow subclusters. (d,e) UMAP visualizations of selected T/NK and monocyte/DC populations in the COMBAT CITE-seq dataset, colored by modality and cell type across the three methods. (f) Quantitative comparison of the same local cross-modality metrics for the selected COMBAT subclusters. The local cross-modality metrics are computed in the existing integrated embedding without reclustering and are defined in Supplementary Section 3. Higher opposite-modality label agreement and hit@k indicate better local alignment of RNA and ADT cells with the same fine-grained annotation.

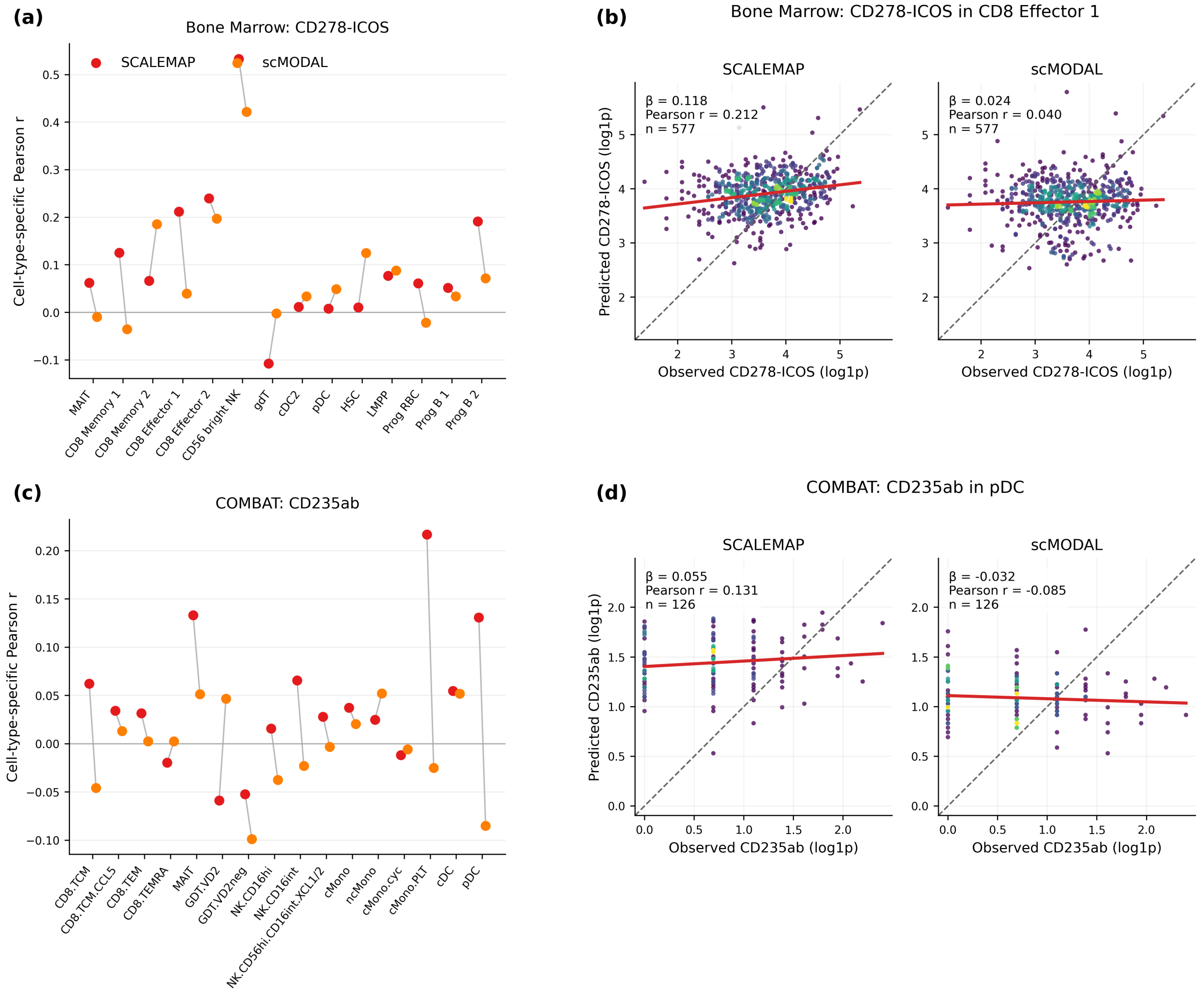


**Fig. S4** Cell-type-level RNA-to-protein prediction in the Bone Marrow and COMBAT datasets. (a) Pearson correlations for CD278-ICOS prediction by SCALEMAP and scMODAL across the Bone Marrow cell types shown in Supplementary Fig. S3. (b) Observed-versus-predicted CD278-ICOS abundance in CD8 Effector 1 cells, the displayed Bone Marrow cell type with the largest between-method Pearson-correlation difference for this marker. (c) Pearson correlations for CD235ab prediction by SCALEMAP and scMODAL across the displayed COMBAT cell types. (d) Observed-versus-predicted CD235ab abundance in pDC cells, the displayed COMBAT cell type with the largest between-method Pearson-correlation difference for this marker. In b and d, the solid red line is the ordinary least-squares regression fit, with its slope reported as $\beta$, and the dashed identity line indicates exact agreement between observed and predicted abundance.

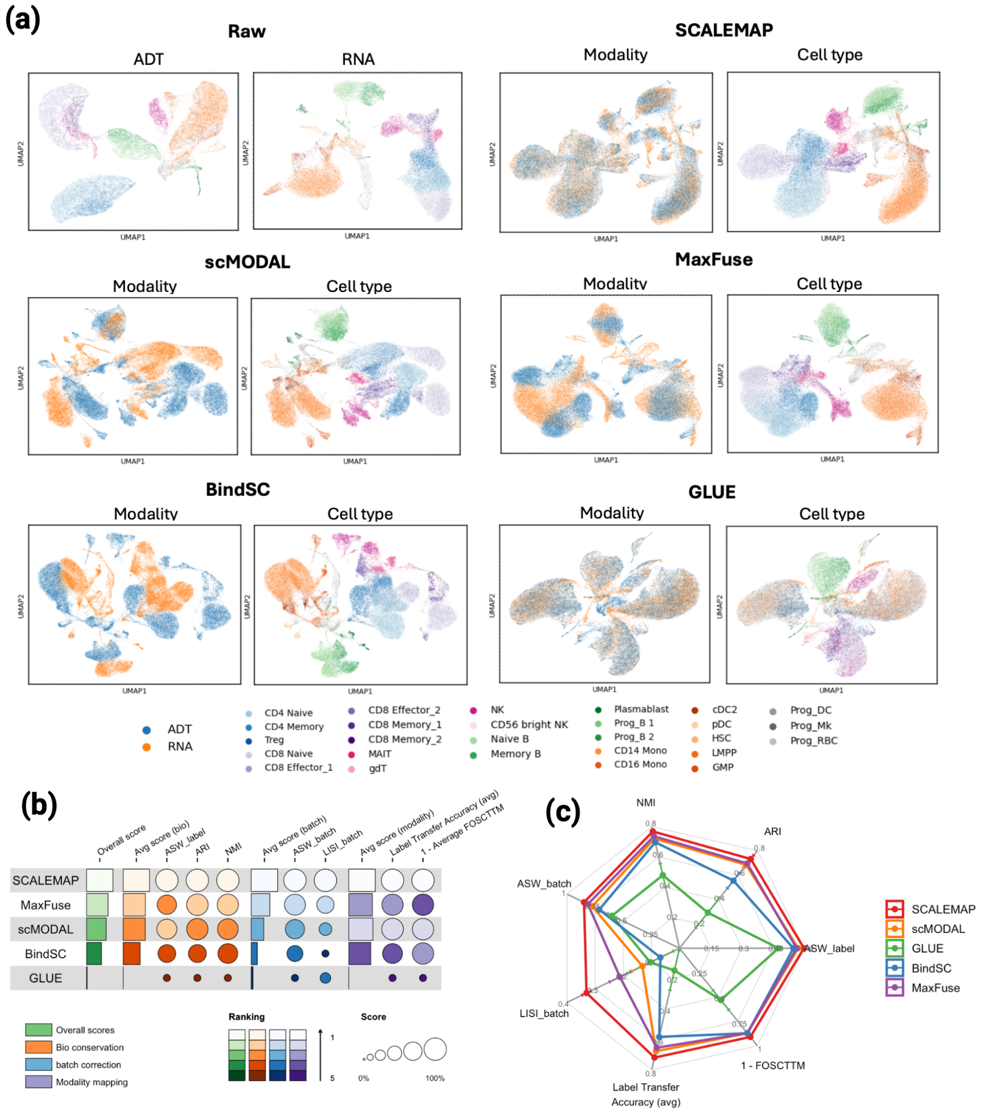


**Fig. S5** Bone Marrow CITE-seq RNA–ADT benchmark.(a) UMAP embeddings before integration and after integration with SCALEMAP, scMODAL, MaxFuse, BindSC and GLUE. Unintegrated embeddings are colored by cell type; integrated embeddings are colored by modality and cell type. (b) Integration scores and method rankings for biological conservation, batch correction and modality alignment. Circle size indicates metric score and color intensity indicates rank, with lighter colors denoting higher ranks. (c) Radar plots of the seven integration metrics defined in Supplementary Section 3; higher values indicate better performance.

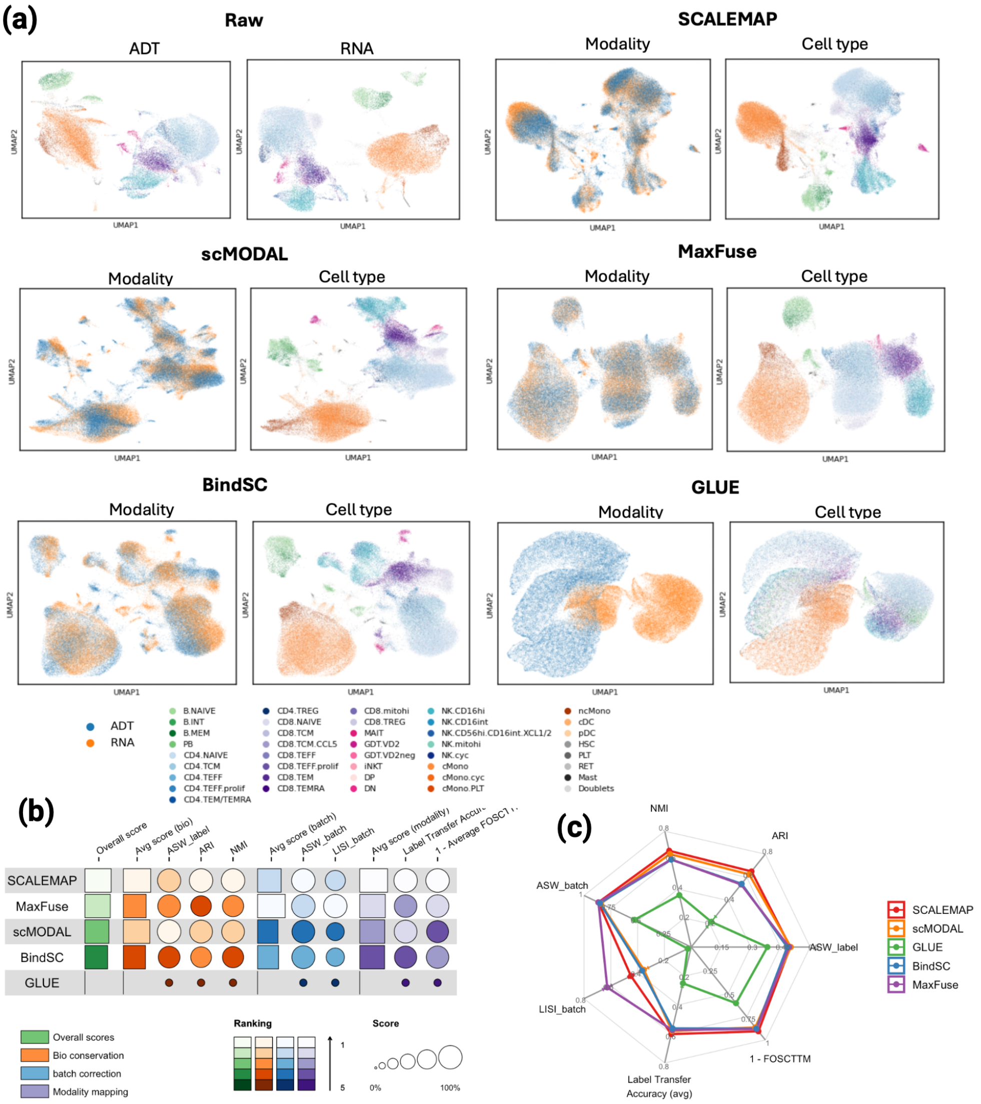


**Fig. S6** COMBAT CITE-seq RNA–ADT benchmark using the 0.05 subset. (a) UMAP embeddings before integration and after integration with SCALEMAP, scMODAL, MaxFuse, BindSC and GLUE. Unintegrated embeddings are colored by cell type; integrated embeddings are colored by modality and cell type. (b) Integration scores and method rankings for biological conservation, batch correction and modality alignment. Circle size indicates metric score and color intensity indicates rank, with lighter colors denoting higher ranks. (c) Radar plots of the seven integration metrics defined in Supplementary Section 3; higher values indicate better performance.

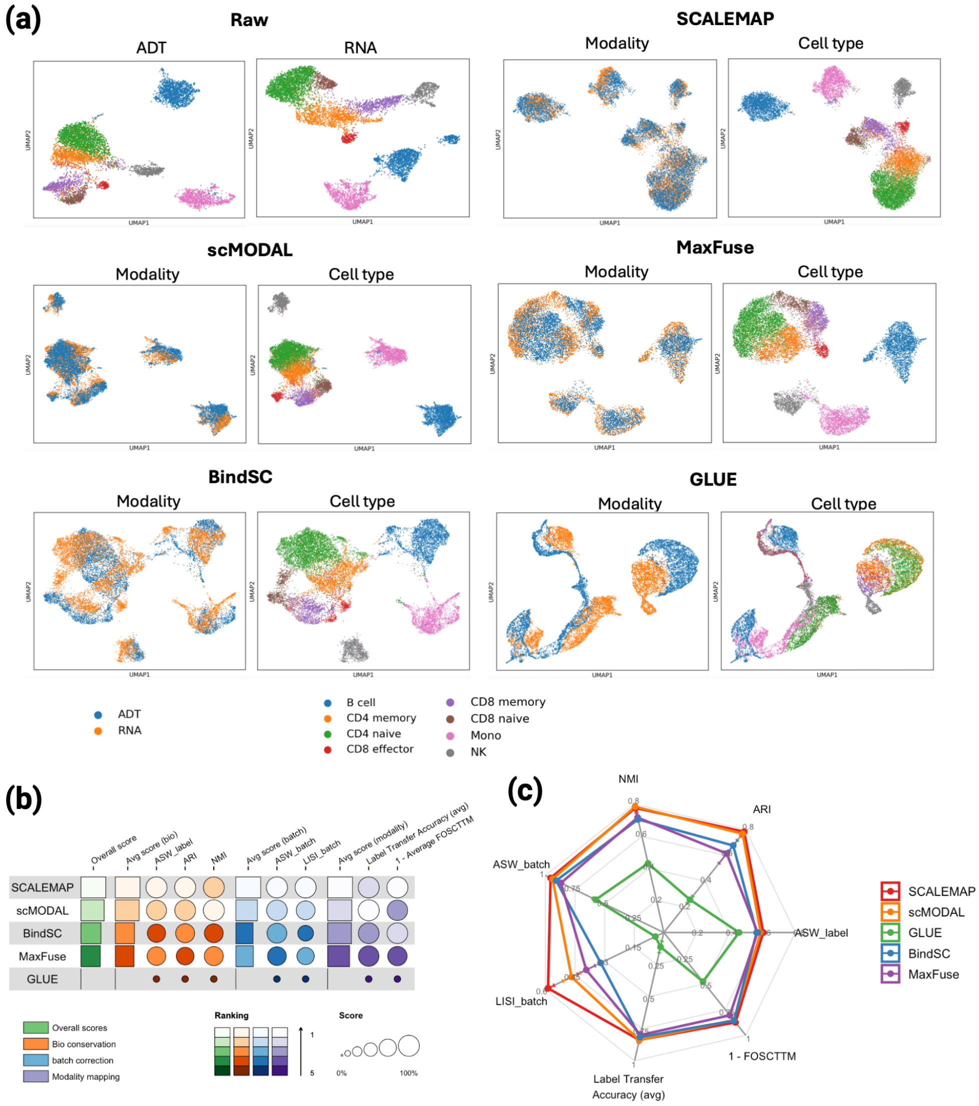


**Fig. S7** PBMC TEA-seq RNA–ADT benchmark. (a) UMAP embeddings before integration and after integration with SCALEMAP, scMODAL, MaxFuse, BindSC and GLUE. Unintegrated embeddings are colored by cell type; integrated embeddings are colored by modality and cell type. (b) Integration scores and method rankings for biological conservation, batch correction and modality alignment. Circle size indicates metric score and color intensity indicates rank, with lighter colors denoting higher ranks. (c) Radar plots of the seven integration metrics defined in Supplementary Section 3; higher values indicate better performance.

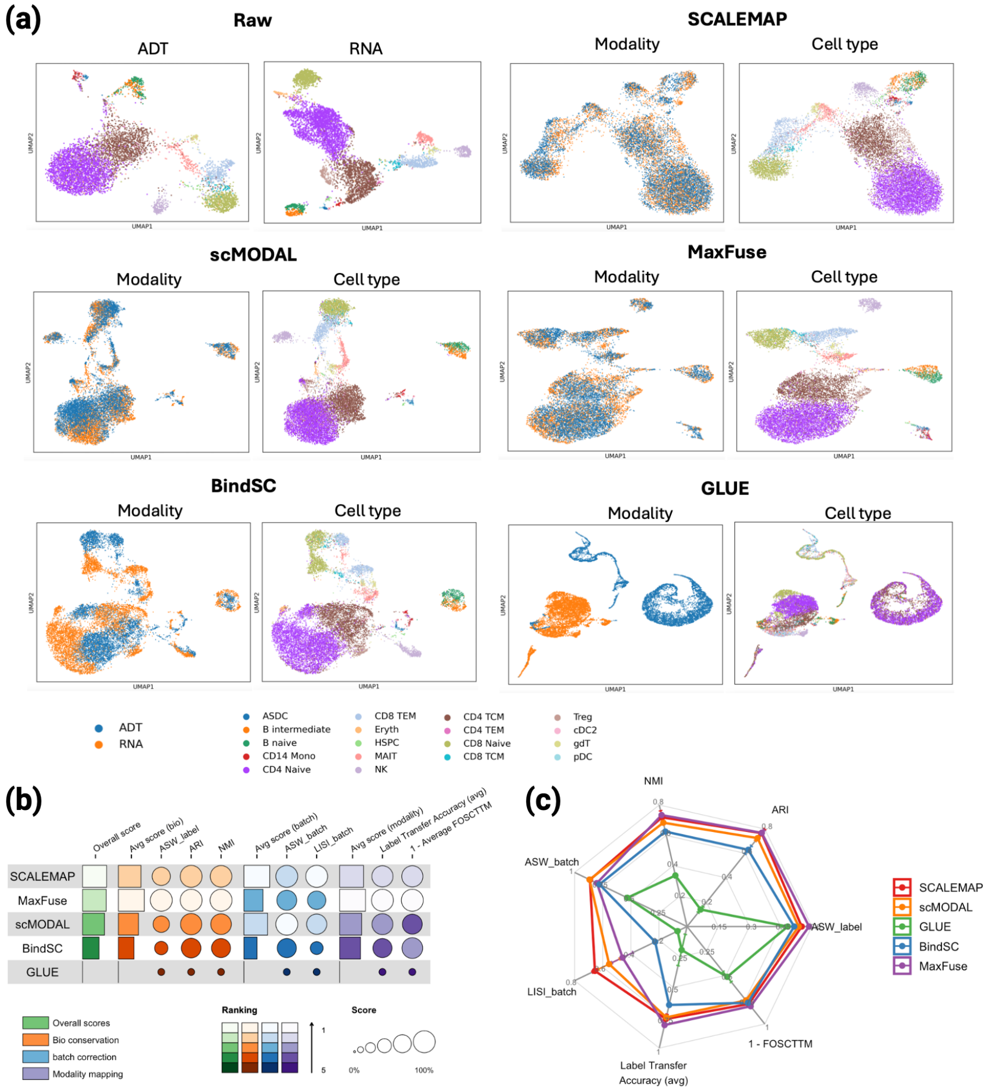


**Fig. S8** PBMC ASAPseqCtrl RNA–ADT benchmark.(a) UMAP embeddings before integration and after integration with SCALEMAP, scMODAL, MaxFuse, BindSC and GLUE. Unintegrated embeddings are colored by cell type; integrated embeddings are colored by modality and cell type. (b) Integration scores and method rankings for biological conservation, batch correction and modality alignment. Circle size indicates metric score and color intensity indicates rank, with lighter colors denoting higher ranks. (c) Radar plots of the seven integration metrics defined in Supplementary Section 3; higher values indicate better performance.

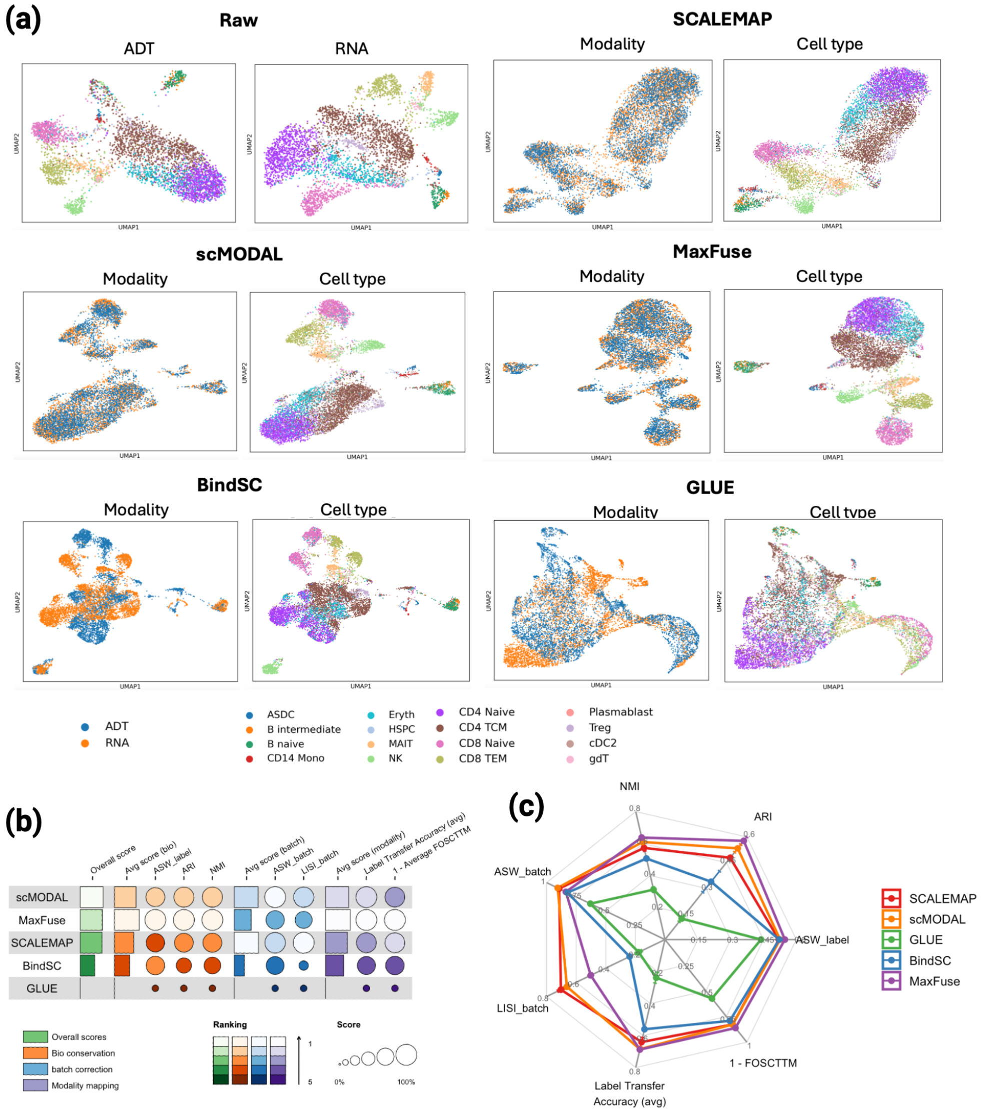


**Fig. S9** PBMC ASAPseqStim RNA–ADT benchmark. (a) UMAP embeddings before integration and after integration with SCALEMAP, scMODAL, MaxFuse, BindSC and GLUE. Unintegrated embeddings are colored by cell type; integrated embeddings are colored by modality and cell type. (b) Integration scores and method rankings for biological conservation, batch correction and modality alignment. Circle size indicates metric score and color intensity indicates rank, with lighter colors denoting higher ranks. (c) Radar plots of the seven integration metrics defined in Supplementary Section 3; higher values indicate better performance.

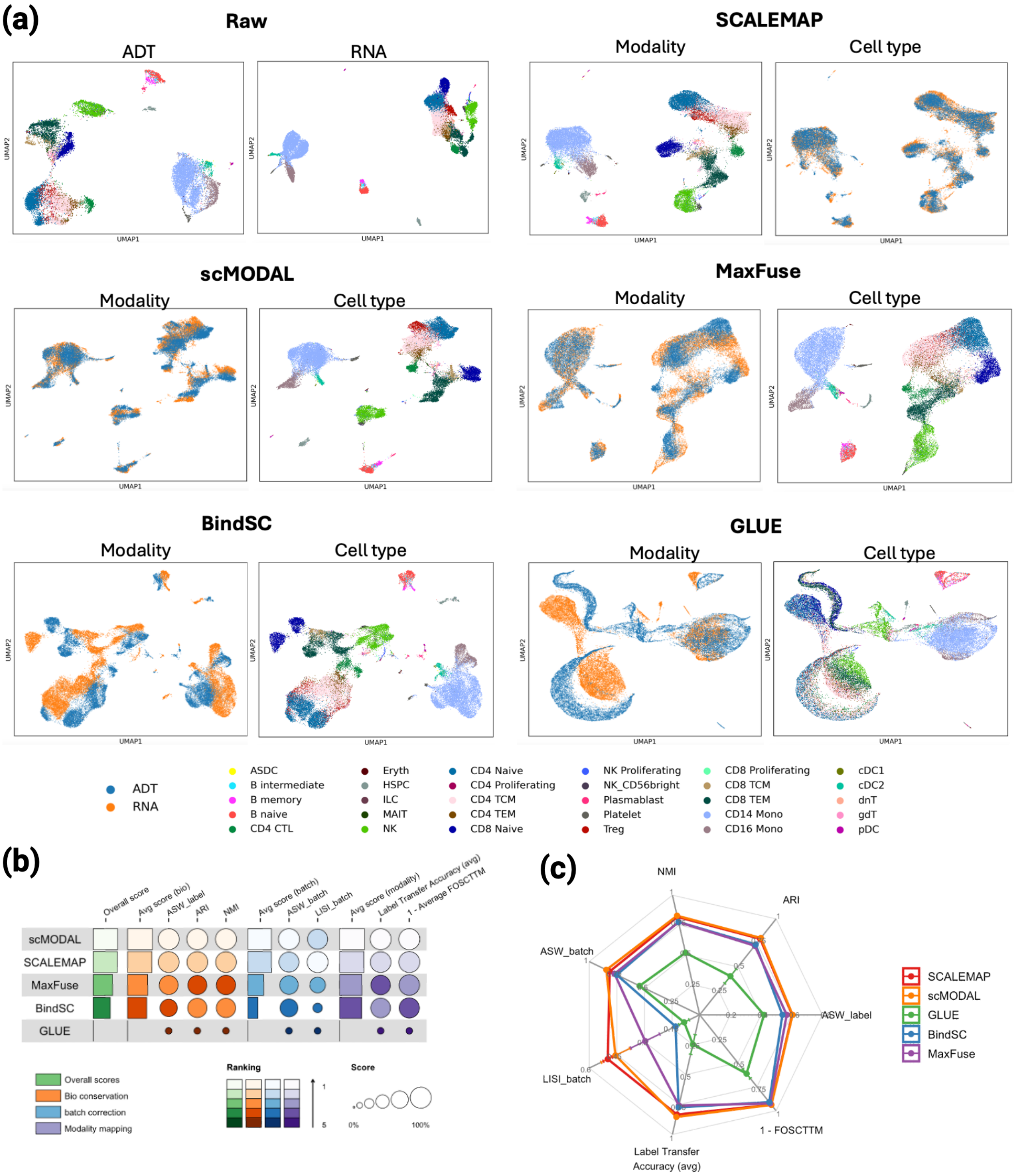


**Fig. S10** PBMC CITE-seq RNA–ADT benchmark. (a) UMAP embeddings before integration and after integration with SCALEMAP, scMODAL, MaxFuse, BindSC and GLUE. Unintegrated embeddings are colored by cell type; integrated embeddings are colored by modality and cell type. (b) Integration scores and method rankings for biological conservation, batch correction and modality alignment. Circle size indicates metric score and color intensity indicates rank, with lighter colors denoting higher ranks. (c) Radar plots of the seven integration metrics defined in Supplementary Section 3; higher values indicate better performance.

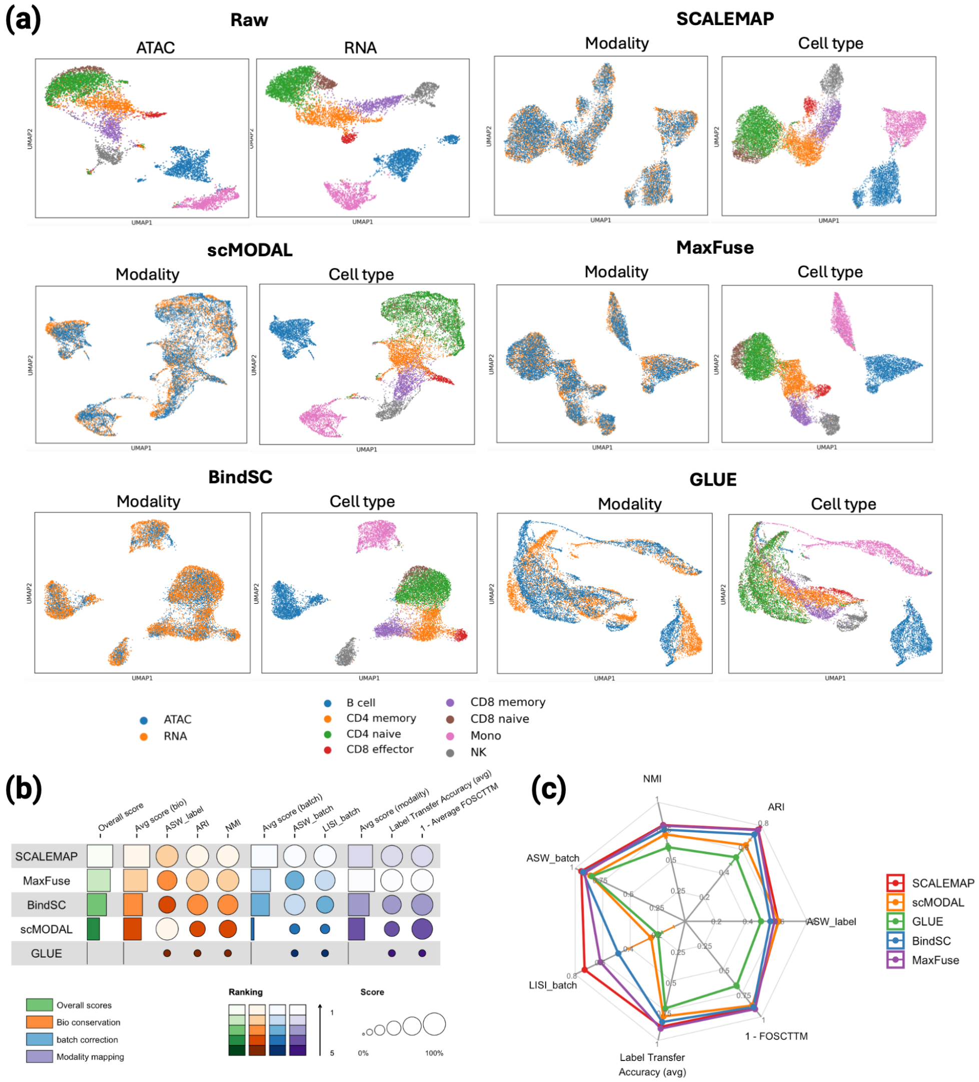


**Fig. S11** PBMC TEA-seq RNA–ATAC benchmark. (a) UMAP embeddings before integration and after integration with SCALEMAP, scMODAL, MaxFuse, BindSC and GLUE. Unintegrated embeddings are colored by cell type; integrated embeddings are colored by modality and cell type. (b) Integration scores and method rankings for biological conservation, batch correction and modality alignment. Circle size indicates metric score and color intensity indicates rank, with lighter colors denoting higher ranks. (c) Radar plots of the seven integration metrics defined in Supplementary Section 3; higher values indicate better performance.

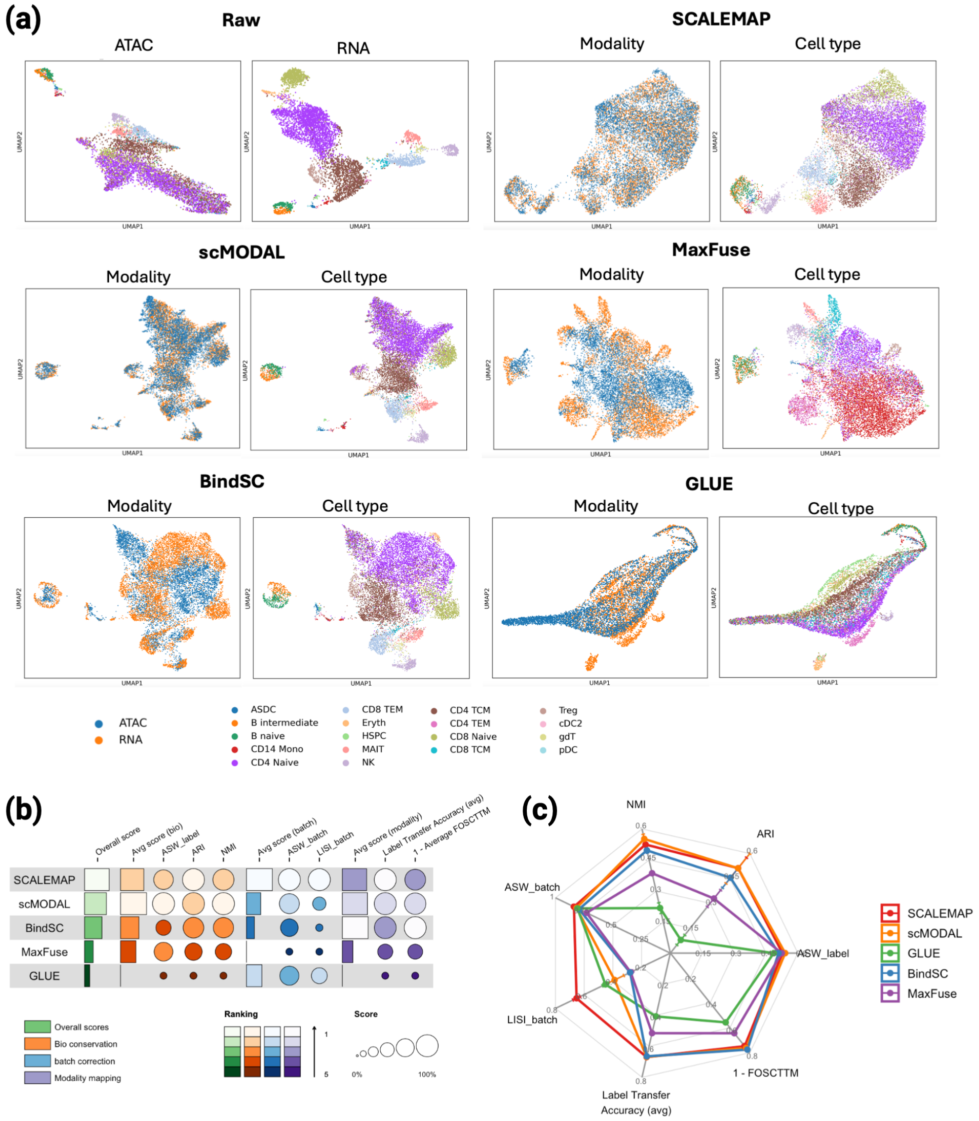


**Fig. S12** PBMC ASAPseqCtrl RNA–ATAC benchmark. (a) UMAP embeddings before integration and after integration with SCALEMAP, scMODAL, MaxFuse, BindSC and GLUE. Unintegrated embeddings are colored by cell type; integrated embeddings are colored by modality and cell type. (b) Integration scores and method rankings for biological conservation, batch correction and modality alignment. Circle size indicates metric score and color intensity indicates rank, with lighter colors denoting higher ranks. (c) Radar plots of the seven integration metrics defined in Supplementary Section 3; higher values indicate better performance.

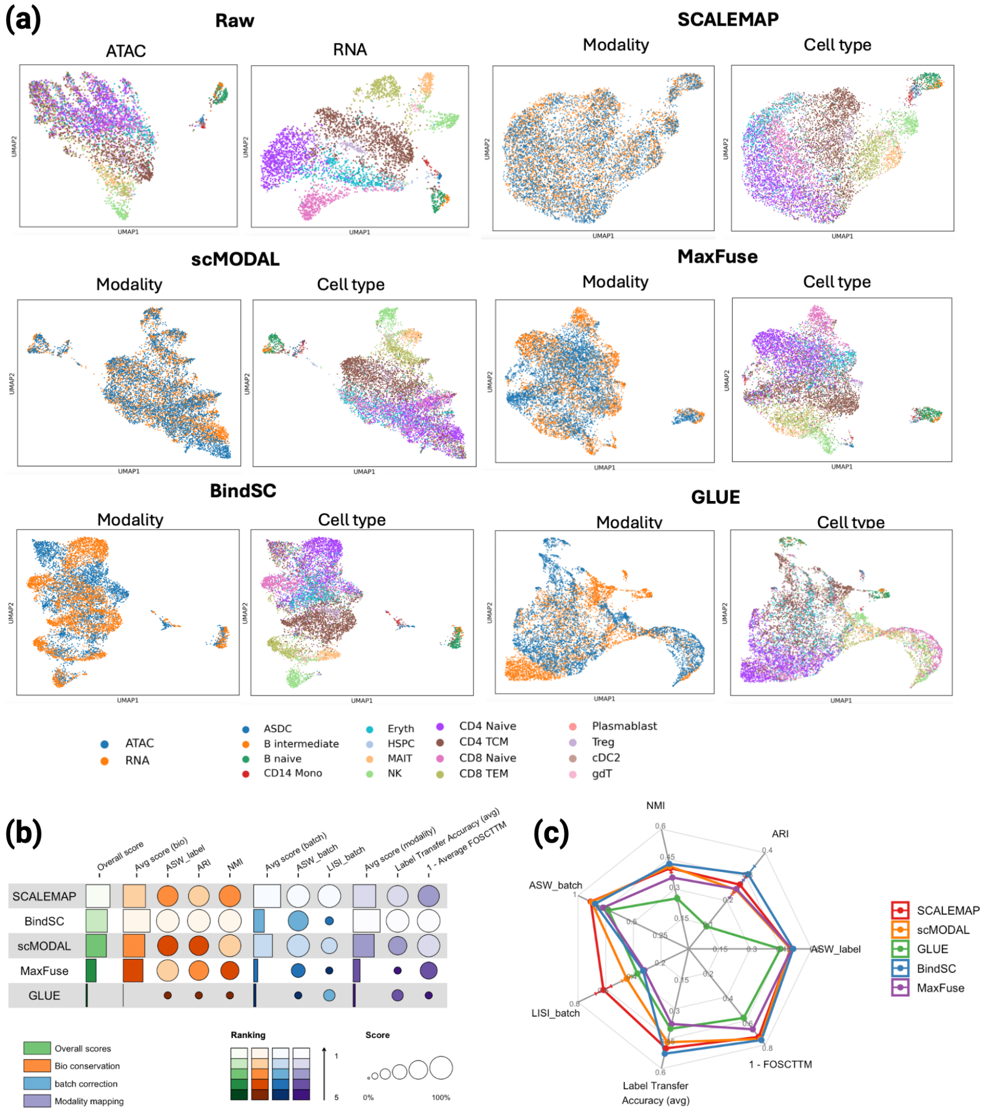


**Fig. S13** PBMC ASAPseqStim RNA–ATAC benchmark. (a) UMAP embeddings before integration and after integration with SCALEMAP, scMODAL, MaxFuse, BindSC and GLUE. Unintegrated embeddings are colored by cell type; integrated embeddings are colored by modality and cell type. (b) Integration scores and method rankings for biological conservation, batch correction and modality alignment. Circle size indicates metric score and color intensity indicates rank, with lighter colors denoting higher ranks. (c) Radar plots of the seven integration metrics defined in Supplementary Section 3; higher values indicate better performance.

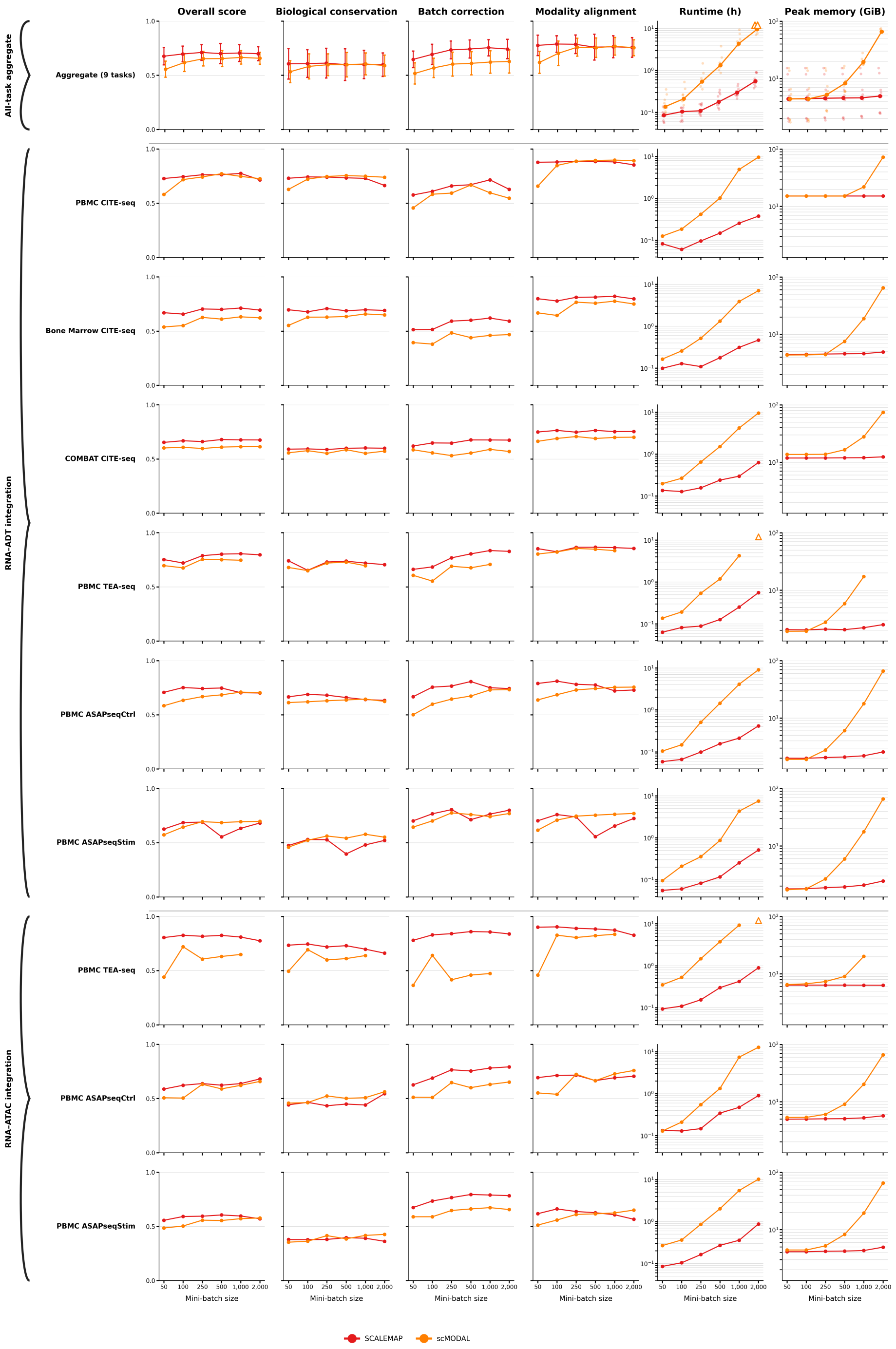


**Fig. S14** Task-level mini-batch sensitivity and computational scaling. Columns show overall integration score, biological conservation, batch correction, modality alignment, full-command runtime in hours and peak resident memory in GiB. The first row summarizes all nine tasks, followed by six RNA–ADT and three RNA–ATAC task rows, grouped by the braces at left. Red denotes SCALEMAP and orange denotes scMODAL. In the aggregate row, performance lines show means and error bars show the sample standard deviation of absolute scores across available tasks. Main-text Fig. 6 uses the sample standard deviation of signed changes from batch size 1,000. Aggregate runtime and memory lines show medians, with faint points indicating individual completed tasks. Task rows show individual results at each batch size. Runtime and memory use logarithmic axes. Open triangles indicate the two scMODAL PBMC TEA-seq tasks that exceeded the 12-h limit at batch size 2,000; unavailable scores and memory values are omitted. All nine SCALEMAP tasks completed at every batch size, whereas scMODAL aggregate summaries at batch size 2,000 comprise seven completed tasks.
